# Kinetic parameter prediction using neural networks identifies limitations to C_4_ photosynthesis

**DOI:** 10.1101/2025.07.16.665120

**Authors:** Philipp Wendering, John Ferguson, Rudan Xu, Johannes Kromdijk, Zoran Nikoloski

## Abstract

Large-scale kinetic models of photosynthesis enable time-resolved predictions of traits related to this key process, and provide the means to identify factors limiting photosynthesis. However, their use is currently limited by the lack of efficient approaches to estimate the hundreds of genotype-specific kinetic parameters. Here, we present C4TUNE, an artificial neural network, which can efficiently predict parameters of a large-scale photosynthesis model from photosynthesis response curves. C4TUNE was trained on a biologically-relevant synthetic dataset comprising matched samples of parameters and response curves obtained using a C_4_ photosynthesis kinetic model. To speed up the training of C4TUNE, we devised a surrogate neural network to predict photosynthesis response curves directly from the model parameters and environmental inputs. Given response curves as input, we showed that over 99% of the parameter vectors predicted by C4TUNE could be used directly in simulation of the kinetic model and resulted in excellent fits. Finally, we applied C4TUNE to predict parameters for a population of 68 maize genotypes across two seasons. The predicted genotype-specific parameters allowed pinpointing factors that limit photosynthetic efficiency, validated using simulations. Therefore, the use of C4TUNE presents a fast and precise approach for parameter prediction based on minimal datasets.

## Introduction

Photosynthesis is a critical determinant of plant production, yet it is still far from its theoretical efficiency ^1^. Photosynthetic efficiency is determined by multiple factors across different scales, from light interception to CO_2_ conversion and partitioning of carbon assimilates ^2^. Much effort has focused on engineering the core carbon metabolism, on minimizing losses of fixed CO_2_ via the photorespiratory pathway ^3^ as well as on the individual or combined manipulation of enzyme expression in the Calvin-Benson cycle, electron transport, carbon transport, and photoprotection ^4^. Despite the success of these experiment-driven efforts, it remains challenging to identify what limits photosynthetic efficiency, particularly under rapidly changing environmental conditions projected for future climate scenarios ^5,6^. Therefore, a paradigm shift in the search for molecular factors that limit photosynthesis, and their manipulation via gene editing tools, is urgently needed to guide photosynthesis improvements and downstream biotechnological applications.

Kinetic models of photosynthesis offer a tractable way to identify molecular factors that limit photosynthesis across different environments. The existing kinetic models of C_3_ ^7–9^ and C_4_ photosynthesis ^10,11^ comprise reactions involved in gas exchange, light absorption, carbon fixation, along with canonical pathways of sucrose and starch synthesis. These models detail the change in metabolite concentrations in terms of reaction fluxes using a system of ordinary differential equations (ODEs). The reaction fluxes are in turn described by the enzyme kinetics; comprising a combination of Michaelis-Menten, mass action, and convenience kinetics as well as empirical algebraic equations ^7–9,11^. The enzyme kinetics relate metabolite and enzyme concentrations to the flux of a particular reaction using enzyme-specific parameters (e.g., Michaelis-Menten constant, K_M_, and maximum reaction rate of an enzyme, V_max_). The kinetic parameters are properties of enzymes and can be manipulated by gene editing of the corresponding enzyme-coding gene or other upstream regulatory components ^12^, providing a tangible means for testing and validating the identified factors limiting photosynthesis.

Data on kinetic parameters from a population of plants along with read-outs of net photosynthesis rate offer the means to identify kinetic parameters strongly correlated to photosynthesis rate in a single or a set of environments. These parameters are candidates for factors that limit photosynthesis, as their manipulation in a specified direction, i.e., decrease for negatively correlated and increase for positively correlated, may be used to increase photosynthesis. However, obtaining data on kinetic parameters spanning all enzymes in photosynthesis-related pathways is an experimentally unfeasible task, due to the time and resources required. Kinetic models of photosynthesis together with data on photosynthesis-related traits can be used to arrive at model parameterizations following, in principle, two directions: (1) parameter estimation, by model fitting procedures, and (2) parameter prediction, by employing artificial intelligence (i.e., machine or deep learning) models to predict parameters, such that photosynthesis-related traits simulated by the kinetic model match the measured traits. Genotype-specific parameterization can in turn be used to identify parameters that limit photosynthesis following the previously described correlation-based procedure.

The proposed direction is made tractable by advances in deep learning for model parameterization ^13^. Despite advances in approaches for parameter estimation ^14–16^, such as those relying on Monte Carlo sampling, parameter estimation methods are time-intensive and their out-of-the-box usage in biology is nontrivial due to the need for modeling skills required. In contrast, recently developed deep learning approaches employ physics-informed neural networks, where the ODE model equations are used to constrain the training process ^17,18^, as well as generative models, i.e., models trained to predict model parameterizations with defined characteristics without any input other than random noise ^19,20^. The proposed generative approaches facilitate the sampling of millions of model parameterizations with stable characteristics that agree with experimental observations in a matter of seconds. For model training, both approaches can also integrate various omics data by pre-processing them into steady-state profiles via thermodynamic stoichiometric modeling to obtain a set of reaction fluxes, metabolite concentrations, and thermodynamic parameters. To generate training data or evaluate model convergence properties using the steady-state profiles, these approaches assume standard enzyme kinetics ^21,22^, not typical for models of photosynthesis (see above). Therefore, the parameter estimation in currently available large-scale photosynthesis models poses additional challenges because they comprise combinations of enzyme kinetics and empirical equations, and due to the sparsity of data that can be used to validate model parameterizations.

Here we developed C4TUNE (C_4_ Tuning Engine), a deep learning approach that predicts complete parameterizations of a C_4_ photosynthesis model from gas exchange data that are routinely measured in photosynthesis research, i.e., response curves of net CO_2_ assimilation rate, *A*_*net*_, to different ambient CO_2_ partial pressures and light intensities. C4TUNE uses synthetic, biologically constrained training data generated by ODE model simulation of sampled parameter vectors. During model training, the biological relevance of the predicted parameters is ensured by: (1) respecting the covariance structure of the model parameters and (2) assessing the simulated *A*_*net*_response curves. To enable time-resolved response curve simulation using the predicted parameters during model training in reasonable time, we used a surrogate model trained to reproduce ODE model simulations. To this end, we built a surrogate model for a large-scale kinetic model of C_4_ photosynthesis ^11^ that comprises 236 tunable parameters for 123 reactions and 109 mass balances. As a result, the trained C4TUNE model efficiently predicts model parameters given *A*_*net*_ measurements as input. We demonstrated that by sampling inputs (with added noise) and subsequent parameter prediction, thousands of possible model parameterizations can be obtained in only few seconds. Finally, we applied C4TUNE to predict parameters for 68 genotypes of a multiple parent advanced generation intercross (MAGIC) maize population for two growing seasons. Using correlation-based analysis, we pinpointed parameters that limit photosynthesis, raising candidates for finetuning of this key process. Therefore, C4TUNE provides the means for efficient and complete model parameterization, which can be used to derive valuable insights into limiting factors and enable simulations of genotype-specific photosynthesis in unseen environments.

## Results

### Efficient prediction of C_4_ photosynthesis parameters in a large-scale kinetic model

The major contribution of our work is a procedure for training an artificial neutral network, C4TUNE, that predicts parameters of a large-scale kinetic model of C_4_ photosynthesis ^11^ given gas exchange measurements. Specifically, C4TUNE makes use of the response of net CO_2_ assimilation rate, *A*_*net*_, to ambient CO_2_ partial pressures and light intensities, gathered in A/CO_2_ and A/light curves, respectively, that are facile to measure ^23^. C4TUNE results in a model parameterization that can be used to accurately simulate these experimental measurements (Fig. 1), via the generation of a synthetic dataset of matched parameter values and A/CO_2_ and A/light curves, necessary for model training.

**Figure 1.**
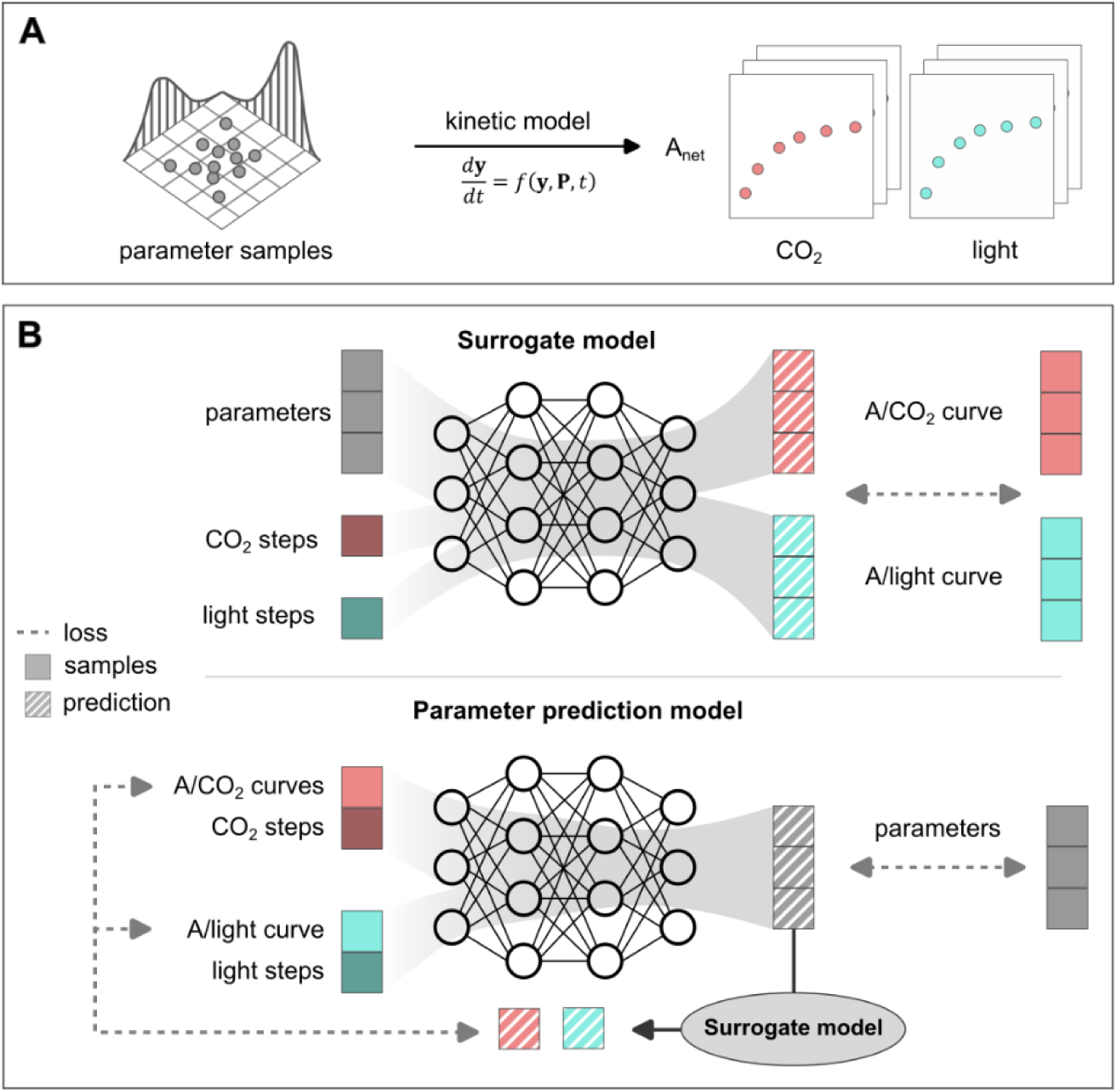
Schematic overview of the generation of artificial training data and training of neural networks in C4TUNE. **A**. Based on a set of initial, feasible parameter sets, parameter samples were drawn from a multi-variate log-normal distribution (n=106). For each of the sampled parameter vectors, responses of the net CO_2_ assimilation rate (*A*_*net*_) to different levels of ambient CO_2_ and light intensity were simulated, resulting in A/CO_2_ and A/light curves, respectively. The simulations were performed using a kinetic model of C_4_ photosynthesis ^11^, where ***y*** denotes the initial state vector of the system of ordinary differential equations (ODEs), ***P*** denotes the vector of parameters, and *t* is the time. **Β.** Inputs and outputs of the trained artificial neural networks. A surrogate model was trained to predict ODE model simulations, *i.e.*, A/CO_2_ and A/light curves based on a parameter vector and CO_2_ and light steps of the *A*_*net*_response curves. The model weights were updated by minimizing a loss function based on the distance between the predicted and samples *A*_*net*_response curves. The parameter prediction model takes A/CO_2_ and A/light curves and the associated CO_2_ and light steps as inputs and predicts a vector of ODE model parameters. The trained surrogate model was in turn used to predict *A*_*net*_ response curves based on the predicted parameter vector. The loss function used to update the weights during the training process of the parameter prediction model was a weighted sum of the distance between the sampled and predicted parameter vectors and the distance between sampled curves and the curves predicted using the surrogate model.

### Generation of a comprehensive synthetic dataset for neural network training

To ensure that C4TUNE can accurately predict model parameters reproducing various shapes of experimentally measured A/CO_2_ and A/light curves, the dataset used for model training must include diverse curve shapes alongside corresponding parameterizations. However, such matched experimental data on *A*_*net*_response curves and model parameters are currently not available. Therefore, we generated synthetic training data by sampling values for each of the 236 tunable parameters (Supplementary Data 1). We then used to simulate A/CO_2_ and A/light curves with the sampled parameter values, thus avoiding the need for matched experimental measurements (Fig. 1A). Whilst the parameter sampling can be performed at random, we opted to make use of genotype-specific parameter values. These were obtained from a Monte-Carlo parameter estimation strategy with data from 68 maize accessions from gas exchange experiments conducted in 2022 and 2023 (n=136, Methods) ^24^ Whilst the estimation of the 236 parameters is not statistically acceptable due to the comparatively low number of measured data points, these estimates provide feasible model parameterizations for the measured *A*_*net*_ response curves. In addition, the parameters result in simulated A/CO_2_ and A/light curves of low mean squared error (MSE) with a median of 2.38 with respect to the measurements.

Next, we sampled 1000 parameter vectors from log-normal distributions for each parameter with mean and standard deviation derived from the initial set of genotype-specific estimates. From these, we found that for 287 parameter vectors, at least one *A*_*net*_response curve simulation was infeasible. We note that a simulation was considered infeasible if the model ODEs could not be integrated or the resulting curve was not biologically relevant (see Methods for precise definition). To increase the probability of sampling feasible parameter vectors, we used the covariance structure of the feasible parameter vectors, determined above, to draw all subsequent random samples from a multi-variate log-normal distribution that resembles the structure of the feasible vectors. As a result, 10^6^ additional random parameter samples were drawn and used to simulate *A*_*net*_ response curves, out of which a smaller fraction (23.6%) were infeasible. A two-dimensional non-linear embedding of the parameter space, including feasible, infeasible, and initial parameter vectors, demonstrates the uniform coverage of the parameter space (Fig. 2A). This observation also emphasizes the difficulty in distinguishing feasible from infeasible parameter vectors.

**Figure 2.**
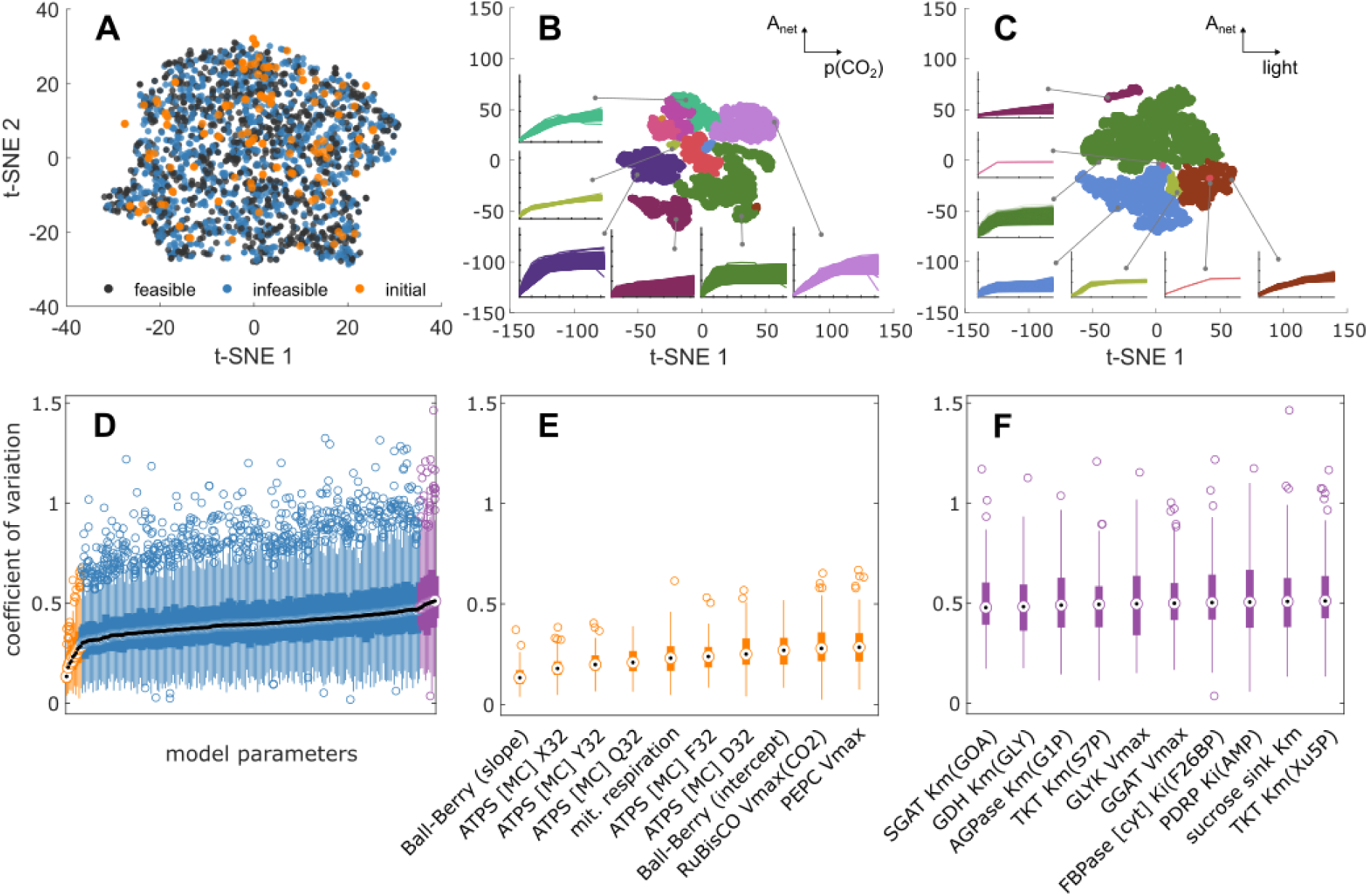
Analysis of the parameter sampling and simulation of response curves. **A.** Non-linear embedding of the sampled parameter vectors using t-SNE ^25^. The embedding was performed on a set composed of randomly selected feasible (n=1000, black) and infeasible (n=1000, blue) as well as the initial parameter sets that were used to estimate the covariance structure (n=136, orange). A parameter vector is feasible if the resulting model can be simulated and the resulting *A*_*net*_ response curves are biologically relevant; otherwise, a parameter vector is considered infeasible (**Methods**). **(B, C)** Non-linear embedding of simulated *A*_*net*_response curves to CO_2_ (B) and light (C) using t-SNE (n=10,000). To obtain clusters of different curve types, spectral embedding was performed based on the t-SNE embedding, followed by K-medoids clustering of the results (K=12 for A/CO_2_ and K=7 for A/light). A full representation of the curves in each cluster is shown in Supplementary Figs. 12 and 13. (D) Distributions of coefficients of variation (CVs) of parameters associated with highly similar A/CO_2_ and A/light curves. A random sample parameter vectors was selected (n=10,000) and the associated *A*_*net*_ response curves were compared with all other curves in the entire dataset to find clusters of similar curves. The CV per parameter was calculated for each cluster that contained at least five members (n=152). (**E, F**)The ten parameters with lowest (orange, E) and highest median CV (purple, F) in ascending order, corresponding to the color code in (D). ATPS: ATP synthase (chloroplast), PEPC: phosphoenolpyruvate carboxylase, SGAT: serine---glyoxylate transaminase, GDH: glycine cleavage system, AGPase: ADP-glucose pyrophosphorylase, TKT: transketolase, GLYK: D-glycerate 3-kinase, GGAT: glycine transaminase, FBPase: fructose-bisphosphatase, PDRP: PPDK regulatory protein, PPDK: pyruvate, phosphate dikinase; parameters of the electron transport equations ^11,44^ X32: light partition coefficient, Y32: J_max_ partition coefficient, Q32: curvature parameter (θ) for the response of electron transport to light, F32: combined absorption, *abs*, and light quality factor, *f*: *abs* ⋅ (1 − *f*), D32: ATP/electron ratio (whole chain+Q-cycle); GOA: glyoxylate, GLY: glycine, G1P: glucose-1-phosphate, S7P: sedoheptulose-7-phosphate, F26BP: fructose-2,6-bisphosphate, Xu5P: xylulose-5-phosphate, MC: mesophyll cell, Km: Michaelis-Menten constant, Vmax: maximum reaction velocity, Ki: inhibitory constant, the specificity of the constants is indicated in parentheses.

In addition to ensuring sufficient sampling coverage of the parameter space, a synthetic dataset should capture the variety of A/CO_2_ and A/light curves shapes that may be obtained from measurements. To investigate whether the synthetic dataset contains distinct classes of curve shapes, we performed clustering for the generated A/CO_2_ and A/light curves separately. To this end, the simulated curves were embedded using t-distributed stochastic neighbor embedding (t-SNE, van der Maaten & Hinton, 2008), followed by spectral clustering using K-medoids as clustering algorithm (K=12 (A/CO_2_), K=7 (A/light), Methods). The numbers of clusters for both curve types were determined by choosing the clustering that a high median Silhouette Index (SI, SI=0.99 (A/CO_2_), SI=0.95 (A/light)), yielding a good separation of the curves, which did not include one giant and several very small clusters. As a result, the clustering revealed groups of distinct curve shapes, which mainly differ by the point where *A*_*net*_ reaches a plateau and the magnitude of the plateau (Fig. 2B,C), supporting the biological relevance of the generated dataset.

To further determine whether parameters are identifiable, i.e., a unique value corresponds to a single pair of A/CO_2_ and A/light curves, we determined the coefficient of variation (CV) per parameter across identical curves. A curve was termed identical if the average distance of *A*_*net*_across all CO_2_ and light steps were below a defined threshold, which was chosen well below the measurement error (see Methods). To identify groups of identical curve pairs, we randomly selected 10,000 curve pairs from the dataset and scanned them for identical curve pairs. In total, we identified 152 groups of identical curve pairs, for which we calculated the CV of the underlying parameters. As a result, we found that the median CV between identical curves ranged between 0.13 and 0.51 with an average of 0.39 over the 236 parameters (Fig. 2D). For instance, the Michaelis-Menten constant of RuBisCO for CO_2_ (median CV=0.36) showed values between 0.005 and 0.017 for a cluster of five identical curves (Supplementary Fig. 1). Overall, the parameters with the lowest median CVs were related to model inputs, e.g., the Ball-Berry model ^26^, parameters related to photosynthetic electron transport rate (e.g., X32, Y32, Q32, F32, and D32), mitochondrial respiration, and the V_max_ values of the RuBisCO carboxylation activity and PEP carboxylase (Fig. 2E). In contrast, parameters with the highest median CV values were related to enzymes in the Calvin-Benson-Bassham (CBB) cycle (transketolase), sucrose synthesis and sink (sucrose sink reaction, fructose-bisphosphatase), C_4_ photosynthetic carbon assimilation cycle (pyruvate, phosphate dikinase (PPDK) regulatory protein), photorespiration (glycine transaminase, D-glycerate 3-kinase, glycine cleavage system, serine-glyoxylate transaminase), and starch synthesis (ADP-glucose pyrophosphorylase) (Fig. 2F). This is in line with the expectation that parameters related to processes that integrate model inputs are more likely to affect *A*_*net*_than those of downstream processes. Taken together, we conclude that the parameters of the model used are not mathematically identifiable in a strict sense, i.e., different parameter sets can result in identical simulation results in terms of the considered A/CO_2_ and A/light curves, within experimental error. However, the maximum CV between identical curves is below 1 for 75% of the parameters, indicating small differences between the values for each parameter that result in identical curves that warrant their prediction from data.

### Neural network as a surrogate for kinetic model simulations

The limited parameter identifiability represents a challenge in training a model to predict parameter values. To address this issue, we included the MSE between measured response curve and the curve simulated with the predicted parameters as an additional term in the loss function for training C4TUNE. However, such a loss function requires millions of kinetic model simulations for corresponding parameterization during the training process. This is practically infeasible given that each model simulation to steady state takes at least 1.3 seconds per curve pair. To remedy this problem, we trained a neural network that predicts the simulation results of the kinetic model; therefore, this neural network acts as a surrogate for the kinetic model simulation. A similar idea underpins a previous study that predicts parameters of Michaelis-Menten kinetics in a model of *E. coli*’s metabolism ^27^. In brief, the developed surrogate model has three different input branches for the parameters and the two *A*_*net*_ response curves including CO_2_ and light inputs. All three inputs are processed to obtain embeddings, whereby the parameter embedding is combined with each of the two curve embeddings via an attention layer. The surrogate model has two output branches, one yielding the A/CO_2_ curve and one for the A/light curve (Fig. 1, Supplementary Fig. 2).

The performance of the trained surrogate model was assessed on an unseen test set comprising 20% of the generated dataset (n=152,960) (Fig. 3A, B). The median MSE was 0.75 ± 0.40 (median ± median absolute deviation (MAD)) with 95% of the MSE values below 3.55. The performance was slightly better for A/CO_2_ curves (Fig. 3A, MSE=0.56 ± 0.41, R^2^=0.99 ± 0.00, median ± MAD), compared to A/light curves (Fig. 3B, MSE=0.84 ± 0.45, R^2^=0.99 ± 0.01, median ± MAD). Moreover, the MSE values from A/CO_2_ and A/light curves generated with the same parameter values showed a Pearson correlation coefficient of 0.78. This indicates, as expected, that for most of the parameter vectors, there is dependence between the prediction qualities for both curve types. Overall, the surrogate model predictions showed a very high similarity to the curves simulated by the ODE model, showing that the model can safely be used to replace the model ODEs for the purpose of curve prediction in model training.

**Figure 3.**
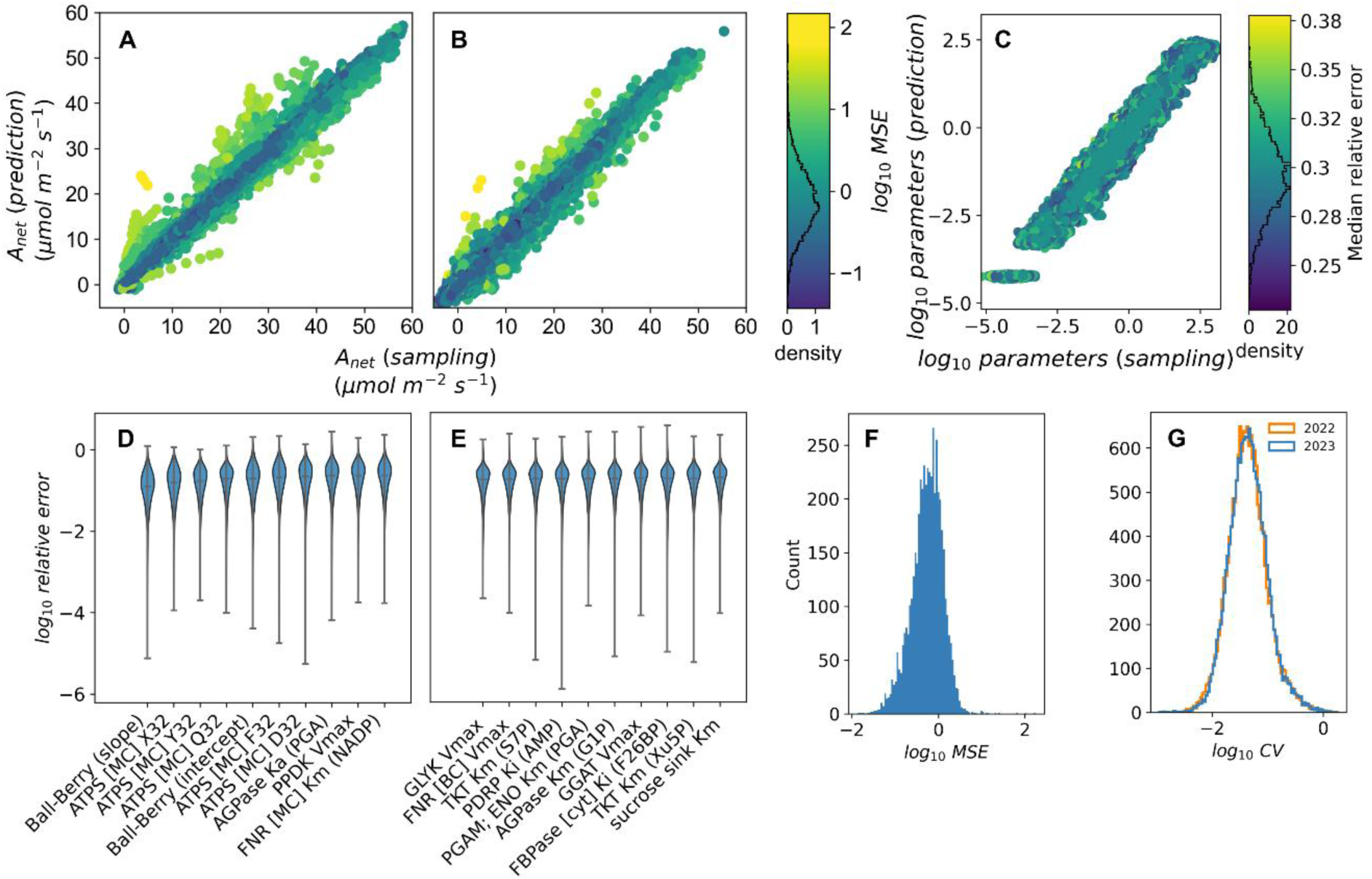
Performance of the trained surrogate and parameter prediction models. **A, B.** Sampled and predicted *A*_*net*_response curves to CO_2_ (A) and light steps (B) using the trained surrogate model, based on 10,000 randomly selected parameter vectors from an unseen test set. The color gradient indicates the average mean squared error (MSE) across both curve types. The black line in the colorbar shows the distribution of the logarithmic average MSE values. **C.** Sampled and predicted parameter values using the trained parameter prediction model (n=5,000). The color gradient indicates the average relative error per parameter vector. The black line in the colorbar shows the distribution of median relative errors. **D, E.** Distributions of logarithmic relative errors of the ten parameters with the lowest (D) and highest median relative errors (E) in ascending order. The whiskers extend to the lowest and highest values, respectively, and the vertical line within the violins represents the median value. **F.** Distribution of logarithmic MSE values of ODE model simulations using the predicted parameters (n=5,000), compared with the curves simulated with the corresponding parameter samples. The MSE values are the average MSE values of both curve types. **G.** Logarithmic coefficients of variation (CV) of predicted parameters for 68 maize accession and year grown in 2022 and 2023, respectively. For each accession and year, ten A/CO_2_ and A/light curves were randomly generated within the respective experimental error and the associated parameters were predicted using **C4TUNE**. ATPS: ATP-synthase (chloroplast), AGPase: ADP-glucose pyrophosphorylase, PPDK: pyruvate, phosphate dikinase, FNR: ferredoxin---NADP+ reductase, GLYK: D-glycerate 3-kinase, TKT: transketolase, PDRP: PPDK regulatory protein, PGAM: phosphoglycerate mutase, ENO: 2-phosphoglycerate enolase, GGAT: glycine transaminase, FBPase: fructose-bisphosphatase; parameters of the electron transport equations ^11,44^ X32: light partition coefficient, Y32: J_max_ partition coefficient, Q32: curvature parameter (θ) for the response of electron transport to light, F32: combined absorption, *abs*, and light quality factor, *f*: *abs* ⋅ (1 − *f*), D32: ATP/electron ratio (whole chain+Q-cycle); PGA: 3-phosphoglycerate, S7P: sedoheptulose-7-phosphate, G1P: glucose-1-phosphate, F26BP: fructose-2,6-bisphosphate, Xu5P: xylulose-5-phosphate, MC: mesophyll cell, Km: Michaelis-Menten constant, Vmax: maximum reaction velocity, Ki: inhibitory constant, the specificity of the constants is indicated in parentheses.

### C4TUNE efficiently predicts model parameterizations that match measured response curves

The architecture of C4TUNE for the prediction of kinetic model parameters was comprised of two input-processing branches for the two curve types and their CO_2_ and light inputs, resulting in embeddings that were concatenated and further processed by dense layers to render the final output, i.e., deviations from the average parameter values (Supplementary Fig. 3). Instead of parameter values, the model predicts deviations from average parameter values that follow the covariance structure of the parameter space. C4TUNE is tuned to optimize a weighted sum of the relative error between true and predicted parameter values and the MSE between predicted curves from the trained surrogate model and the true values (Methods). The performance of the final, trained model was assessed using a random sub-sample of the unseen test set comprising 5,000 samples, resulting in a median relative error between true and predicted parameter values of 0.30 ± 0.16 (median ± MAD, Fig. 3C). A closer inspection of the distribution of relative errors of predicted parameters revealed that the median relative errors across the test samples showed differences across the parameters with the highest median relative error of 0.40 ± 0.20 (median ± MAD) observed for the K_M_ value of the sucrose sink reaction for sucrose and the lowest median relative error of 0.13 ± 0.07 observed for the slope of the Ball-Berry model. In agreement with the observed CV values of the parameters across identical curves (Pearson r=0.96), the parameters with the highest and lowest relative errors largely overlapped with the parameters with high and low CV, respectively (Fig. 3D,E). This shows that the predictability of the parameters is directly related to their unique association with distinct curve shapes influencing the capacity of the neural network to learn the relationship between parameter values and associated *A*_*net*_ response curves.

Further, the median MSE between true A/CO_2_ and A/light curves and the curves predicted by the surrogate model was 0.06 ± 0.03 (median ± MAD). Notably, this error is more than ten-fold lower than the MSE of the surrogate model with sampled parameters as input. On one hand, this shows that the predicted parameters result in surrogate model curve predictions which are practically indistinguishable from the samples curves. On the other hand, it may also indicate that C4TUNE is overfitted to predict parameters that result in exceptional curve predictions with the surrogate model, but not necessarily when using them to integrate the model ODEs. Given that the surrogate model does not discriminate between feasible and infeasible parameter sets, the low MSE between predicted and true curves for the parameter prediction model does not guarantee that C4TUNE results in feasible parameter vectors for ODE model simulation. To test whether the predicted parameters can be used to simulate the ODE model with a similar performance to the surrogate model, we re-ran the curve simulations using the ODE model with 5,000 randomly selected predicted parameter vectors. We found that over 99% of the predicted parameter vectors resulted in successful simulations, with a median MSE of 0.58 ± 0.30 (median ± MAD) with 95% of the MSE values lower than 1.86 (Fig. 3F). Here, too, the median MSE for A/light curves (0.64 ± 0.29) was higher than for A/CO_2_ curves (0.48 ± 0.29). Importantly, most predicted parameters could be used in simulations using the original ODE model with a low error, showing that C4TUNE is not overfitted to the surrogate model.

Taken together, the presented parameter prediction model showed excellent performance in recovering ODE model parameters underlying a diverse set of simulated *A*_*net*_ response curves. The results demonstrated the applicability of the approach to produce results that can be used for downstream analysis, such as additional curve simulations for unseen conditions or simulation of mutant phenotypes. Finally, the limited identifiability of the parameters with respect to the considered A/CO_2_ and A/light curves was reflected in a lower bound on the relative errors between predicted and true parameter values. We expect that a large proportion of the limited identifiability and resulting reduction in predictability is caused by compensatory effect between parameters, especially V_max_ and K_M_ values of the same enzyme. However, the identifiability of some of the parameters could likely be improved by including additional experiments, e.g., gas exchange measurements under different environmental conditions, protein abundances, or metabolite concentrations.

### C4TUNE provides precise maize genotype-specific model parameters that result in low simulation error

To evaluate the performance of C4TUNE when applied to experimental measurements, we predicted parameters for 68 maize accessions grown in 2022 and 2023. To verify that the predicted parameters are indeed relevant, we performed simulations for *A*_*net*_response curves with the surrogate and ODE model for both growing seasons, respectively. The median MSE values between the experimental measurements and surrogate model predictions were 2.26 (2022) and 1.25 (2023). For the ODE model simulations, the median MSE values were 2.38 and 1.43 for 2022 and 2023, respectively. This result shows that, while the measured A/CO_2_ and A/light curve pairs were not considered alongside the training response curves generated by the ODE model, C4TUNE was still able to predict parameters that resulted in curves closely resembling the experimental observations (Supplementary Fig. 4).

The results presented above were obtained by predicting parameter vectors for average values of *A*_*net*_response curves over the biological replicates. However, there can be considerable variation across replicates, with median CV values of 0.14 ± 0.05 (2022) and 0.16 ± 0.06 (2023) for A/CO_2_ curves and 0.20 ± 0.10 (2022) and 0.23 ± 0.10 (2023) for A/light curves. To investigate how the uncertainty in experimental measurements is reflected in the predicted parameters, we sampled ten curves from a normal distribution with mean and standard deviation specific to the measurement for each accession and year. Next, we used C4TUNE to predict parameters which were in turn used for simulation with the surrogate model and the ODE model. The resulting curves showed median MSE values of 3.09 for 2022 and 5.33 for 2023 using the surrogate model (median *R*^2^ = 0.98 and 0.97), and 4.73 and 7.95, respectively, using the ODE model (median *R*^2^ = 0.98 and 0.95, Supplementary Fig. 5). The median CV of the predicted parameters was 0.04 for both years (Figure 3G), demonstrating that C4TUNE can reliably predict parameters for perturbed curve inputs, sampled within the experimental error. These values were almost ten-fold lower compared to the CVs of parameters associated with identical curves, despite the higher variation in the synthetic response curves used for training. This observation showed that the prediction performance of C4TUNE is robust against experimental uncertainty in the input data. It further indicated that the parameter identifiability is higher based on the maize-specific *A*_*net*_ response curves. Given the higher CV of parameters associated with identical curves in the dataset, this also means that C4TUNE is less likely to include extreme values of parameter values that result in equally accurate curve simulations.

### Predicted model parameters pinpoint photosynthetic limitations

Next, we used the accession-specific model parameterizations for the 2022 growing season to identify parameters that are highly correlated with changes in net assimilation rate. To this end, we determined the pairwise correlations between the predicted parameters and *A*_*net*_ values at each CO_2_ or light intensity step. In addition, the pairwise Pearson correlations, *r*, between *A*_*net*_ values at different steps of the A/CO_2_ and A/light curves were calculated to assess their agreement. We found that the *A*_*net*_ values at the different CO_2_ levels were highly correlated with each other (0.68 ≤ min(|*r*|) ≤ 0.88), with pronounced, strong correlations between 25 µbar and 400 µbar and from 600 µbar to 1250 µbar (Fig. 4**Figure 2**A). For the A/light curves, *A*_*net*_values measured at high light intensities were highly correlated across the genotypes, while there were low and even partly negative correlations between *A*_*net*_values measured at low and high light intensities (0.01 ≤ min(|*r*|) ≤ 0.36, Fig. 4**Figure 2**B), indicating pronounced genotype-by-environment interactions for different CO_2_ inputs (as environments). Given that the model structure and initial concentrations remained the same across genotypes, the photosynthesis parameters are expected to determine the observed dependencies between the steps of the *A*_*net*_ response curves as they allow the ODE simulations to match genotype-specific curve pairs. Therefore, the minimum absolute correlations per CO_2_ or light step between the measured *A*_*net*_values were used as thresholds to determine representative associations between *A*_*net*_ response curves and model parameters.

**Figure 4.**
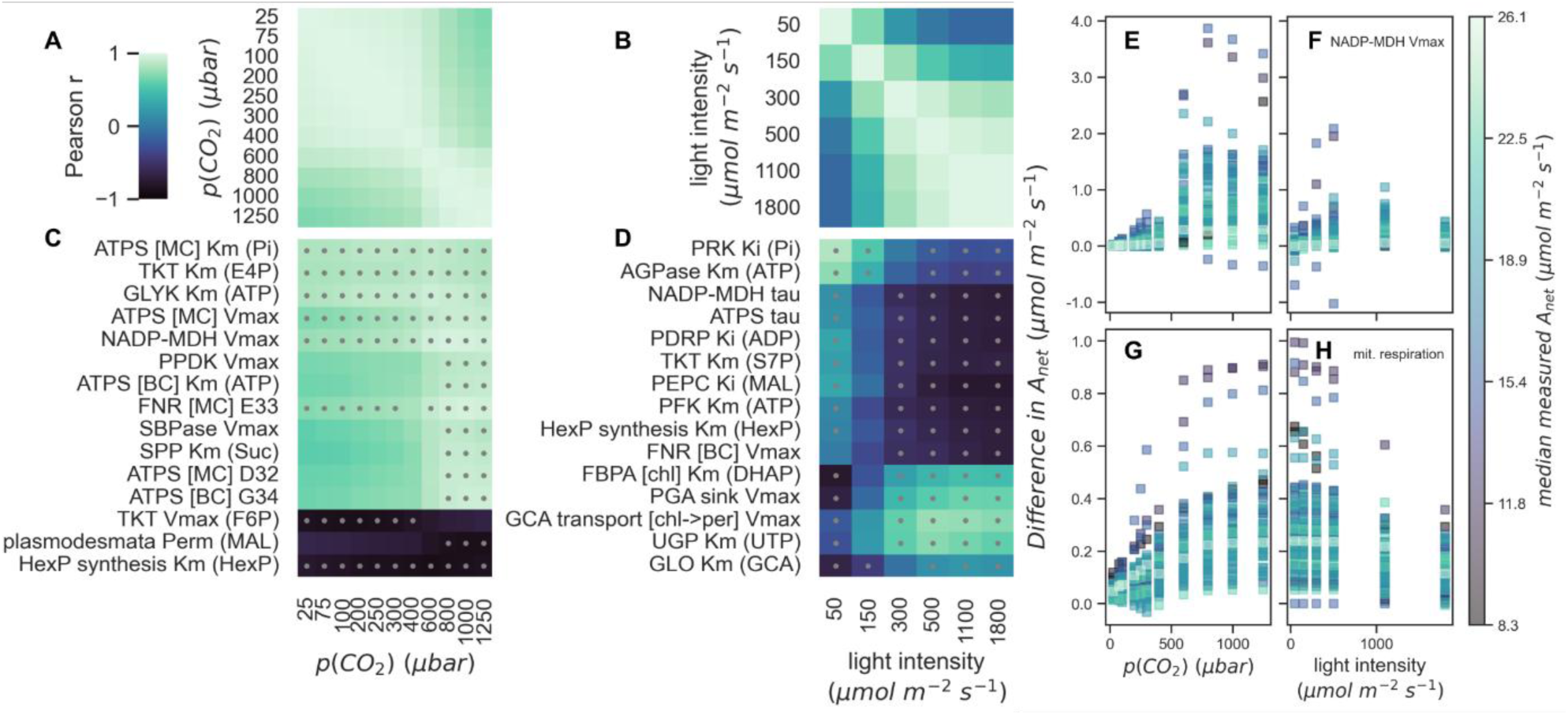
Correlations between *A*_*net*_ and parameter values. Pairwise Pearson correlations between **A., B.** *A*_*net*_values at different CO_2_ and light steps and **C., D.** between *A*_*net*_values and log-transformed parameter values were determined. The *A*_*net*_response curves (A/CO_2_ and A/light curves) were obtained from 68 maize genotypes grown in 2022. The heatmaps in panels C and D show the correlations of the 15 parameters that ranked highest when sorted by the median correlation across all CO_2_ or light steps, respectively. The color code indicating correlation is the same across all panels as indicated by the common color bar. Grey dots in the heatmap tiles indicate that the respective correlation coefficient is higher than the minimum of pairwise correlation between A_net_ values measured at the different ambient CO_2_ or light intensity levels, respectively (shown in panels A and B). Parameters that showed the highest sign-consistent correlations to A/CO_2_ and A/light were considered as potential targets for improvement of photosynthetic efficiency. Their values were updated individually to the highest or lowest value of the parameter predicted across the entire population, depending on the sign of the correlation. The updated parameter sets were used to simulate A/CO_2_ and A/light curves using the ODE model and the differences to the original curve were quantified for all 68 genotypes. The simulated curves with the highest median differences resulted from increased V_max_ values of the NADP-malate dehydrogenase enzyme, which showed strong correlations to A/CO_2_ curves (**E, F)**, and from reduced mitochondrial respiration, which showed a strong correlation to A/light curves (**G, H**). The colors in panels E-H indicate the median *A*_*net*_ for each curve type per accession, as shown in the colorbar. ATPS: ATP synthase, TKT: transketolase, GLYK: D-glycerate 3-kinase, SPP: sucrose-phosphate phosphatase, SBPase: sedoheptulose-bisphosphatase, NADPH-MDH: malate dehydrogenase, PPDK: pyruvate, phosphate dikinase, PPDK_I: inactive PPDK, FNR: ferredoxin---NADP+ reductase, RuBisCO: ribulose-1,5-bisphosphate carboxylase/oxygenase, UGP: UTP---glucose-1-phosphate uridylyltransferase, MEP: proton:pyruvate cotransporter, PDRP: PPDK regulatory protein, GLO: glycolate oxidase, FBPA: fructose-bisphosphate aldolase, AGPase: ADP-glucose pyrophosphorylase, PRK: phosphoribulokinase, PFK: 6-phosphofructo-2-kinase, PEPC: phosphoenolpyruvate carboxylase, E4P: erythrose-4-phosphate, Suc: sucrose, GAP: glyceraldehyde-3-phosphate, F6P: fructose-6-phosphate, HexP: hexose phoshates, G1P: glucose-1-phosphate, MAL: malate, PYR: pyruvate, GCA: glycolate, PGA: 3-phosphoglycerate, DHAP: dihydroxyacetone-phosphate, S7P: sedoheptulose-7-phosphate:, Km: Michaelis-Menten constant, Jmax: maximum rate of photosynthetic electron transport, Vmax: maximal reaction rate, Perm: permeability, tau: rate constant of light-regulated enzyme activation, D32: ATP/electron ratio (whole chain+Q-cycle), G34: ATP/electron ratio (whole chain), E33: NADPH/electron ratio (whole chain+Q-cycle), chl: chloroplast, per: peroxisome.

We found that some of the parameters whose values showed strong association with *A*_*net*_ also exhibited sign-consistent correlations across the CO_2_ and light steps of the two curve types (Fig. 4C). Moreover, we found that the correlation of the parameters with *A*_*net*_ was dependent on the CO_2_ or light level for most of the parameters. For instance, the permeability of plasmodesmata for malate or the V_max_ value of PPDK only showed high correlations with *A*_*net*_ at high ambient CO_2_ partial pressures. This trend was even stronger for the parameters with highest correlations with A/light curves (Fig. 4D). Here, we also identified parameters where correlations with *A*_*net*_changed signs across different light intensities, that may be relevant in a canopy setting ^28^. For instance, this was observed for the K_M_ value of the chloroplastic fructose-bisphosphate aldolase for dihydroxyacetone-phosphate, which suggests that increasing its value leads to a higher photosynthetic efficiency at high light intensities, whilst resulting in a decreased efficiency at low light intensity. The opposite effect was observed for the K_i_ value of PEPC for malate: a high value (low inhibitory effect) would be beneficial to photosynthesis at low light intensity but tends to lead to lower *A*_*net*_ at high light intensities, which is relevant with regards to ongoing work to elucidate PEPC structure and function^29–31^. However, parameters showing such ambivalent effects may not be the most suitable targets for increasing photosynthetic efficiency. Therefore, we focused only on parameters exhibiting sign-consistent correlations.

To this end, the obtained correlations were filtered for sign-consistency and magnitude using the thresholds for minimum pairwise correlations determined above. For the response of *A*_*net*_ to ambient CO_2_, we found highly correlated parameters associated with the CBB-cycle (K_M_ value of transketolase for erythrose 4-phosphate), photorespiration (K_M_ values of glycerate kinase for ATP), light reactions (V_max_ and K_M_ of the chloroplastic ATP synthase for phosphate), C_4_ photosynthetic carbon assimilation cycle (V_max_ of NADP malate dehydrogenase), and the generation of hexose phosphates. The parameters with the highest median correlations pertained to reaction steps involved in mitochondrial respiration, C_4_ photosynthetic carbon assimilation cycle, light reactions, transport of C_4_ carboxylic acid and triose phosphate, photorespiration, and sucrose synthesis (Supplementary Data 2). These parameters present possible targets for alleviating limitations on photosynthetic efficiency, due to their consistent and strong associations with *A*_*net*_over multiple CO_2_ partial pressures and light intensities for the considered maize accessions.

To estimate the potential impact of optimizing the identified parameters to improve photosynthetic efficiency, we individually modified their values to the respective most extreme value predicted for the population in 2022. These were then used to parameterize the ODE model and to simulate updated A/CO_2_ and A/light curves. We observed that most of the single parameter changes only resulted in small changes in *A*_*net*_(Supplementary Data 2). However, we found that an increase in the V_max_ of NADP malate dehydrogenase or the V_max_ value of the mesophyll chloroplast ATP synthase were predicted to result in *A*_*net*_ changes between −1.03 and 3.87 µmol m^-2^ s^-1^ (−89% and +83%) across all genotypes and simulated conditions (Fig. 4E,F, Supplementary Data 2). Similarly, our predictions indicated that a decrease in mitochondrial respiration rate, a fixed value that increases cytosolic CO_2_ concentrations in both cell types, or the K_M_ value of ferredoxin-NADP+ reductase for NADPH resulted in changes of *A*_*net*_ in the range from −0.03 to 1.00 µmol m^-2^ s^-1^ (−1477% and 1242%, Fig. 4G,H). The changes in *A*_*net*_were strongest at high CO_2_ levels and low to medium light intensities. Moreover, the parameter changes had the strongest impact on accessions with overall low to medium *A*_*net*_ values (see Supplementary Figs. 6-8 for example curves). Notably, the parameter values were optimized using extreme values within the predicted genotype-specific parameters, while the corresponding extreme parameter values in the entire generated dataset were mostly considerably higher or lower (Supplementary Data 2). Although these could yield greater increases in simulated photosynthetic rates (Supplementary Fig. 9), we confined our optimization to the genotype-specific predictions to prioritize biological relevance of the proposed changes. As a control, we also simulated the increase of RuBisCO content, which has previously been shown to increase light-saturated *A*_*net*_ by 15% in maize ^32^. In these simulations, we increased the V_max_ values of the RuBisCO carboxylation and oxygenation reactions to the respective maximum of the predicted values for this parameter in the employed maize population. As a result, we found that the median increase in the simulated *A*_*net*_ values across all genotypes were in the range from 0.4% to 5.6% (Supplementary Fig. 10). When the maximum values from the artificial dataset were used instead, the increases ranged between - 2.3% and 3.4%.

These results demonstrate that C4TUNE applied to data sets from a population of genotypes can pinpoint enzyme parameters that limit photosynthetic efficiency by performing simple correlation analysis. Furthermore, the correlations across the full range of the A/CO_2_ and A/light curves reveal the impact of the parameter change under different environmental conditions, which could partly be confirmed by model simulations. Therefore, C4TUNE can contribute to the discovery of novel targets for alleviating limitations on photosynthetic efficiency using precise parameter predictions.

## Discussion

Here we present C4TUNE, a deep learning framework for the prediction of C_4_ photosynthesis parameters from gas exchange measurements. The model takes as inputs the environment parameters and *A*_*net*_ measurement of A/CO_2_ and A/light curves and predicts a genotype-specific set of ODE model parameters ^11^. We trained C4TUNE on a large synthetic dataset generated by ODE model simulations from randomly sampled model parameterizations. Using a deep learning architecture to predict model parameters provides multiple advantages over classical approaches: (1) the non-linearities of the ODE model are automatically captured by non-linear activations, (2) additional constraints can be easily integrated into the model and training process, (3) very large datasets can be processed efficiently, and (4) the number of predicted parameters is not limited by the number of measurements as in statistical estimation relying on measures like *χ*^2^ values. For instance, we integrated the known covariance structure of the parameters to transform predicted deviations from average parameter values. To consider the limited identifiability of the model parameters, a second neural network (i.e., surrogate model) was trained to predict A/CO_2_ and A/light curve pairs from environmental parameters and ODE model parameters. This allowed the consideration of parameter simulation errors directly and efficiently during model training. Notably, the known biologically meaningful ranges of *A*_*net*_values as well as constraints on the shapes of predicted simulated A/CO_2_ and A/light curves were integrated via the surrogate model architecture and training. By allowing the prediction of full model parameterizations from A/CO_2_ and A/light curves, C4TUNE increases the number of predicted parameters, compared to previous studies ^8,9,33–36^, facilitating the use of kinetic photosynthesis models in different genotypes.

While neural networks have already been used for kinetic model parameterization ^19,20,27,37^, their application with existing photosynthesis models is not straightforward. The challenges include a high number of parameters and the use of non-standard enzyme kinetics and algebraic equations in the ODEs, which hamper the application of the existing neural-network-based approaches. To address these issues, C4TUNE relies on a surrogate model to evaluate whether the predicted parameters can be used to obtain meaningful model simulations. Whilst the use of a surrogate model has been proposed previously ^27^, here we advanced on this work by also considering the parameter prediction error to train C4TUNE. Importantly, the use of these two neural networks enables parameter prediction independent of the nature of enzyme kinetic formulations used in the ODE model, a strategy that can be easily transferred to predict parameters different large-scale kinetic models.

After model training, the surrogate model achieved a median MSE of 0.75, while classical machine learning approaches like a K-nearest neighbors (K-NN) regression model (K=100) and random forest regression achieved median MSE values of 18.61 and 8.95 (Supplementary Fig. 11A). The trained C4TUNE model predicted model parameters with a median relative error of 0.30 (Supplementary Fig. 11B). While a similar relative error could be achieved by a K-NN (K=100, relative error: 0.30) and a random forest regression model (relative error: 0.30), importantly, parameter predictions from each of these approaches gave rise to high errors in the ODE model simulations. Here, the parameters predicted by C4TUNE resulted in a median MSE of 0.58 (median *R*^2^ = 0.94), while the K-NN and random forest models predictions resulted in two-orders of magnitude higher median MSE values of 72.81 and 84.66 (median *R*^2^ = 0.31 and 0.26), respectively (n=100, Supplementary Fig. 11C,D). These results demonstrate the proposed deep learning approach learned to predict precise and biologically relevant parameter sets, which was not possible using classical machine learning approaches.

C4TUNE was trained to predict parameters using a specific experimental setup. We assumed the availability of both A/CO_2_ and A/light curves, measured at specific ambient CO_2_ partial pressures and light intensities as well as different constant light intensities and CO_2_ levels, respectively. While *A*_*net*_responses to different CO_2_ and light steps within each response curve could be accommodated by imputing *A*_*net*_ values at the currently used steps from a curve fit, the model training would have to be repeated if the curves were measured at different constant light or CO_2_ levels. This includes re-running the automated parameter sampling and either re-training the model from the beginning or starting the training process from the current weights and biases. The sampling process is the most critical step, which can be completed within 12-24 h, depending on the level of parallelization. The subsequent model training takes about 12 h using a standard laptop computer, which is expected to be strongly reduced when the training is started from the current model parameters.

To demonstrate the use of C4TUNE with experimental data, we used it to predict parameters from gas exchange measurements for a population of 68 genotypes from a maize MAGIC population. To assess the quality of the parameter predictions, they were used to predict and simulate the same A/CO_2_ and A/light curves with the surrogate and the ODE model. Ideally, the obtained curve simulations closely match the input curves, indicating that an appropriate model parameterization was identified. Indeed, the obtained curves showed median MSE values of 2.26 and 1.25 using the surrogate model and 2.38 and 1.43 using the ODE model for 2022 and 2023, respectively. This demonstrates that C4TUNE is able to predict parameters for the model of C_4_ photosynthesis by Wang et al., 2021, which result in simulations that closely match the experimental measurements. By correlating the predicted genotype-specific parameter vectors to the measured curve pairs, we identified parameters that limit photosynthesis under the considered environmental conditions. Notably, C4TUNE enables the prediction of all 236 tunable model parameters, vastly increasing the number of parameter predictions and estimations compared to previous studies ^8,9,33–36^. Among these, we identified a set of parameters that showed consistently high correlations with *A*_*net*_ across both curve types, which present potential targets for improving photosynthetic efficiency in the considered accessions. These targets could be partly confirmed by ODE model simulations with optimized parameters derived from within the population of maize genotypes. The simulation results obtained by increasing the V_max_ values of the RuBisCO carboxylation and oxygenation reactions demonstrate that the improvement of photosynthetic efficiency by updating a single factor can be reproduced using the predicted parameters. However, the impact of increasing this parameter was not predicted to improve *A*_*net*_ across all genotypes and conditions equally, indicating that the impact of parameter optimizations varies even between closely related genotypes. This highlights the need for genotype-specific model parameterizations for the identification of targets for improving photosynthetic efficiency.

C4TUNE can be readily applied to populations of different species with C_4_ photosynthesis. This is rendered possible by the facile retraining of C4TUNE within 24 h for a different experimental setup or different C_4_ subtype. For instance, C4TUNE can be used to generate genome-specific kinetic models as additional features in genomic prediction of photosynthesis rate ^24^. Therefore, we expect C4TUNE to facilitate the engineering of and selection for traits linked to photosynthetic efficiency, contributing to an improved photosynthetic efficiency and climate resilience of C_4_ crop species.

## Methods

### C_4_ kinetic model

We made use of a previously published C_4_ photosynthesis ODE model ^11^ with minor modifications ^24^. The changes to the original model include updates to the equilibrium constants, *K*_*eq*_, which were inferred from thermodynamic data using the Equilibrator calculation tool (https://equilibrator.weizmann.ac.il, Beber *et al.*, 2022). Moreover, all K_M_ and K_i_ values of the RuBisCO enzyme were unified between the carboxylation and oxygenation reactions, which were previously distinct between the two reactions in the model. This was done to ensure that only one set of kinetic constants was estimated for RuBisCO ^24^.

### Simulations of *A*_*net*_ response curves

All simulated response curves of *A*_*net*_ to changes in ambient CO_2_ or light intensity were performed while keeping constant all other environmental conditions accounted for in the model. More precisely, we set the ambient temperature to 25 °C, wind speed was set to 3.5 m s^-1^, and relative humidity was set to 65%. The steps in ambient CO_2_ partial pressure for curve simulations were 400, 600, 800, 1000, 1250, 400, 300, 250, 200, 100, 75, 25 µbar at a constant light intensity of 1800 µmol photons m^-2^ s^-1^. For the simulations of light response curves, we used light intensities of 1800, 1100, 500, 300, 150, 50 µmol photons m^-2^ s^-1^ at a constant ambient CO_2_ partial pressure of 400 µbar. The simulation time for the first and sixth timestep was set to 3600 s to guarantee establishment of a steady state in *A*_*net*_ at 400 µbar, while all other simulation intervals were 120 s long. Timepoint six was discarded after the simulation. Moreover, two stop criteria were used in the ODE solver: (1) when *A*_*net*_ reaches a steady state, and (2) when the simulation time on the machine exceeded 30 s. The second criterion was added to exclude models with low convergence times as well as to reduce the time required for individual simulations. These settings are equivalent to the ones used to estimate parameters for maize accessions in a parallel manuscript ^24^. All simulations were carried using Matlab’s *ode15s* solver ^39^ with a relative tolerance of 10^-4^ and the requirement for all state variables to be non-negative.

### Criteria for physiologically relevant model parameterizations

The following criteria were employed to classify kinetic models as relevant: (1) convergence of the ODE solver in the simulation of *A*_*net*_ response curves to both CO_2_ and light intensity, (2) any imaginary part of the simulated curve values does not exceed 10^−9^, (3) the simulated curve must contain at least one positive value, (4) all simulated values are between −10 µmol m^-2^ s^-1^ and 70 µmol m^-2^ s^-1^, and (5) no more than one change in monotonicity per response curve (only one change from increasing to decreasing direction or *vice versa* is allowed).

### Sampling the parameter space of the C_4_ photosynthesis model

The space of parameters associated with physiologically relevant kinetic models is multidimensional and shaped by non-linear relationships between parameters ^19,27^. We assessed the model’s sensitivity to perturbations of the initial parameter values ^11^ and found that many of the resulting kinetic models were infeasible or resulted in physiologically irrelevant *A*_*net*_response curves. Therefore, we relied on previous work, where parameters were estimated for 68 maize (*Zea mays* L.) accessions over two years (Supplementary Note 3, ^24^). Based on these parameter estimates, ***P***_***acc***_, 1000 random parameter samples, ***P***\*_1_, were drawn from a log-normal distribution:

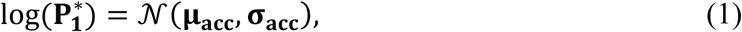

where ***μ***_***acc***_ and ***σ***_***acc***_ are the average and standard deviation of the log-transformed values in ***P***_***acc***_over all accessions. In total, 713 of the 1000 parameter samples resulted in physiologically relevant kinetic models. Using dimensionality reduction approaches, such as principal component analysis (PCA) or t-distributed neighbor embedding (t-SNE) ^25^ it was not possible to distinguish between parameter vectors associated with relevant or irrelevant kinetic models. In a previous study, iterative application of rules derived from decision trees has been used to distinguish between relevant and irrelevant parameter vectors ^40^. While the potential of employing such a rule-based approach is an interesting avenue for future developments, we decided to use a more simplistic approach for this study, which assumes that the joint distribution of the relevant parameter can be captured by their covariance structure. To potentially increase the fraction of relevant parameter vectors, we used the covariance matrix ***∑*** of the log-transformed relevant samples 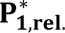 to generate 10^6^ additional parameter samples that follow the same multivariate distribution:

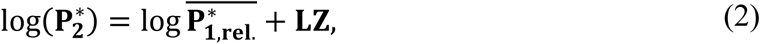

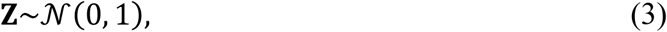

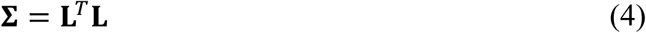

The matrix ***L*** was obtained using Cholesky decomposition of ***∑*** (Eq. (4)). By the multiplication of ***L*** and ***Z*** in Eq. (2), it is ensured that the resulting parameter samples 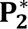 have the mean and standard deviation as well as preserved covariance structure as the samples in 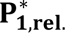. Out of the additional 10^6^ samples, 764,091 were associated with physiologically relevant models, based on the criteria defined above. Compared to the first 1000 samples, the fraction of relevant parameter vectors increased from 71.3% to 76.4%. In total, we obtained 764,804 relevant samples for further analysis.

### Clustering of A/CO_2_ and A/light curves

To assess the diversity of simulated *A*_*net*_ response curves, a clustering analysis was performed on a random subset of 10,000 samples drawn from the generated dataset. For both curves types, the *A*_*net*_values were first embedded using t-SNE ^25^, followed by spectral clustering, using K-medoids as the clustering algorithm based on Euclidean distance. The clustering quality was determined for different values for the number of clusters, 2 ≤ *K* ≤ 15, based on the median Silhouette Index (Supplementary Fig. 12). For both curve types, the optimal *K* was chosen as cluster number that resulted in a good visual separation, while yielding a Silhouette Index, which is close to the optimum for this curve type (*K* = 12 for A/CO_2_ curves and *K* = 7 for A/light curves, Supplementary Figs. 13 and 14).

### Assessment of parameter identifiability

To quantify the identifiability of the parameter space, we determined the variability among parameter vectors that result in identical *A*_*net*_response curves. To this end, we randomly selected 10,000 parameter sets as seeds and proceeded to identify curves in the complete dataset, which had a very low overall Canberra distance to the curves associated with the seed parameter vectors. The distance *d*_*ij*_ between two curves *i* and *j* was defined as

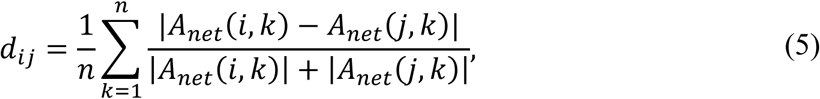

where *n* denotes the number of CO_2_ or light steps used to generate the curve. For each seed, we considered all response curves as identical if the average distance (Eq. (5)) of the A/CO_2_ and A/light curves was below 0.01. For each cluster of identical curves, the coefficient of variation was calculated if the cluster size was greater or equal to five (n=152).

We then determined the coefficient of variation (CV) across the parameter vectors associated with identical curves found for each of the seeds. Seeds for which we identified fewer than ten identical curves were not considered. The CV is defined by

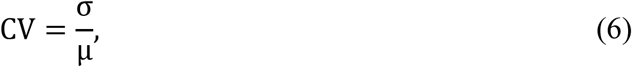

where *μ* and σ denote the mean and standard deviation of the parameter values across a group of identical curve pairs.

### Neural network training

By inspection of the sampled kinetic parameters and associated *A*_*net*_ response curves, we found considerable variation among the parameters associated with identical curves (“Assessment of parameter identifiability”). In a machine learning setting, such many-to-one relations in the training data can be problematic and can hamper the learning process. Therefore, we reasoned that not only the distance between predicted parameter values, but also the curves resulting from the predicted parameters should enter the loss function guiding neural network training for parameter prediction. This arrangement ensures that the model does not only learn to predict the underlying parameters of the response curves, but also considers redundancy in the parameter space.

Given that time needed for simulation of millions of response curves using the ODE solver is not feasible for this model, we trained a surrogate model that predicts *A*_*net*_response curves from the CO_2_ and light intensity inputs and a set of parameters (Fig. 1B). A similar workflow involving a surrogate model has been proposed previously and has been applied to Michaelis-Menten kinetics and a *E. coli* growth model ^27^. The main differences with the workflow by Borisyak and colleagues and the present work include the use of different error functions, as well as our consideration of the error to the associated parameter vector. Due to non-standardized kinetic formulations in the C_4_ photosynthesis model, including algebraic equations, the convergence of the model could not be evaluated automatically. Therefore, the additional error term considering the parameter vector in the loss function was necessary because the use of the surrogate model alone was insufficient to guarantee the prediction of parameter vectors that result in feasible model simulations.

### Surrogate model

The architecture of the surrogate model is composed of three separate input branches processing the input for CO_2_ and light response curves, as well as the parameter vector (n=236). The parameter input is processed by three dense layers of dimension 128, each followed by the rectified linear unit (ReLU) activation function. The inputs relating to the experimental conditions are comprised of the steps in CO_2_ partial pressure (*n*_*co*2_ = 11) or light intensity (*n*_*light*_ = 6), combined with a fixed value of light intensity and CO_2_ partial pressure for the two curves, respectively. The experimental condition inputs are each processed by a single-layer long short-term memory (LSTM), which considers the parameter encoding in each step. The dimension of the hidden layer in both LSTMs is 64. The encodings of experimental conditions and parameters are weighted by an attention mechanism, resulting in a joined encoding with dimension 128. The joined encoding is then combined with each of the encodings of the experimental condition, yielding two dense layers with input dimension 192 and output dimensions corresponding to the size of CO_2_ and light intensity steps, respectively. The output from both final layers is transformed by

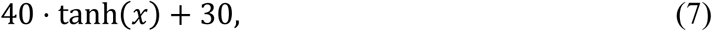

which ensured that the range of model outputs corresponds to the criteria defined above (“Criteria for physiologically relevant model parameterizations”). A depiction of the model architecture can be found in Supplementary Fig. 2.

The surrogate model inputs are z-transformed. For the parameter input, the mean and standard deviation are determined based on the training set and test set, respectively, to avoid data leakage. The means and standard deviations of the CO_2_ partial pressures and light intensity values were calculated from the respective ranges given in section “Simulations of *A*_*net*_response curves”. The model was trained by minimizing a loss function, which combines both the mean squared error (MSE) to the known response curves, but also punishes predictions with more than one change in monotonicity (“Criteria for physiologically relevant model parameterizations”):

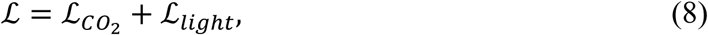

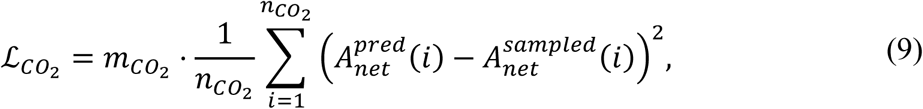

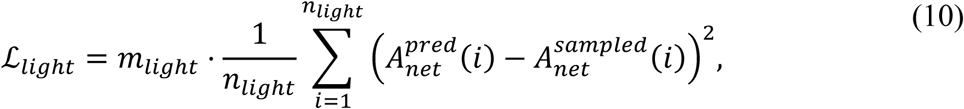

where ℒ denotes the loss function, *m*_*co*2_ and *m*_*light*_ denote the number of changes in monotonicity in the different response curves. The numbers of CO_2_ and light steps are denoted by *n*_*co*_ and *n*_*light*_. The predicted and sampled *A*_*net*_ values are denoted by 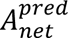.and 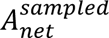, respectively.

Training was performed with an initial learning rate of *α* = 0.027, which decays linearly to 0.01*α* over 15 epochs. Every 15 epochs, the linear schedule was reset, but the initial learning rate was modified by a factor of *γ* = 0.5. The model parameters were updated using stochastic gradient descend with a momentum of 0.78 and Nesterov momentum. Additionally, gradient clipping was applied with a clipping norm of 2.39. Training parameters were optimized by randomized grid search with Adaptive Successive Halving using ray tune 2.37.0 ^41^. In this approach, multiple training runs are performed with parameter combinations randomly drawn from specified parameter distributions or choices. The process is made more efficient by stopping the training runs that perform worse than others with respect to the average loss calculated over the test set. The final model was trained over 60 epochs with a batch size of 8.

### Parameter prediction model (C4TUNE)

The neural network for parameter prediction does not predict the parameter values directly. Instead, it predicts deviations from the average parameter values 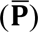 of the training or test set, respectively. The network has two separate input branches, which take the concatenated *A*_*net*_values, CO_2_ or light steps, and respective constant light intensity and CO_2_ partial pressures as inputs. These inputs, of dimensions 23 and 13, respectively, are passed to a single-layer LSTM with a hidden layer dimension of 128. Both LSTMs are followed by a dense layer of dimension 128 and the two encodings are concatenated and passed into a dense layer of dimension 64. This combined curve encoding is then passed through three dense layers with dimensions 128, 256, and 236. The first two of these dense layers are followed by the ReLU activation function while the output of the final layer is transformed by

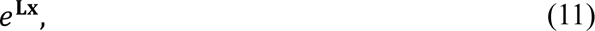

Where ***L*** is again the matrix obtained from applying Cholesky decomposition to the covariance matrix of the log-transformed parameter values of the training or test set (as in Eq. (4)). The final parameter predictions are then calculated by

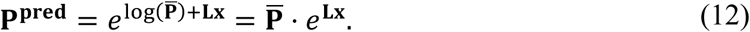

A depiction of the model architecture can be found in Supplementary Fig. 3.

The model was training by minimizing a loss function that combines the relative error between the predicted and known parameter values, and the MSE between *A*_*net*_ response curves predicted using the surrogate model and the known *A*_*net*_ values:

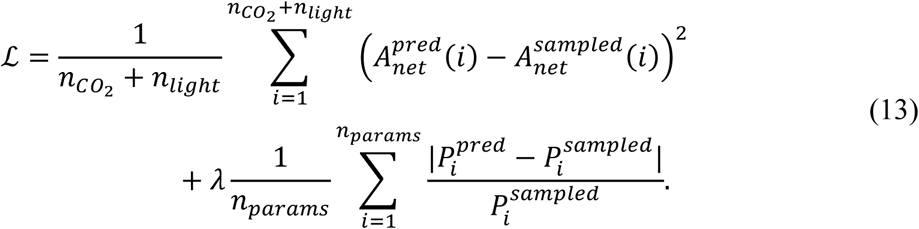

The hyperparameter *λ* balances the two parts of the loss function. Here a value of 9.12 for *λ* was found to yield good results. Moreover, *n*_*params*_ denotes the number of predicted parameters This model was trained with an initial learning rate of *α* = 0.01, and a linear decay to 0.05*α* over 10 epochs. After every 10 epochs, the initial learning rate was multiplied by *γ* = 0.5. The optimization of model parameters was done using stochastic gradient descend with a momentum of 0.70. Further, gradient clipping was performed with a clipping norm of 2.48. The model architecture and training hyperparameters were tuned as described above (“Surrogate model”). The final model was trained for 30 epochs with a batch size of 8.

The neural networks were implemented using Python 3.10.14 using the PyTorch library version 2.5.1 ^42^.

## Data availability

The gas exchange measurements for maize genotypes are available at https://doi.org/10.5281/zenodo.15966533. Part of these data has been used in another study linking photosynthesis-related traits and hyperspectral reflectance data ^43^. The generated artificial data set for neural network training is available at https://doi.org/10.5281/zenodo.15926601.

## Code availability

Custom code for the generation of the artificial dataset as well as code for neural model definition and training are available at https://github.com/pwendering/C4TUNE. This repository also contains the predicted parameters for the maize genotypes.

## Funding

This work was funded by a research grant from the Biotechnology and Biological Sciences Research Council (BB/Y51388X/1) to JK.

## Author contributions

P.W.: Conceptualization, Methodology, Software, Investigation, Writing - Original Draft, Writing - Review & Editing, Visualization. J.F.: Investigation, Data curation. R.X.: Software, Investigation. J.K.: Conceptualization, Writing - Review & Editing, Supervision, Project administration, Funding acquisition. Z.N. Conceptualization, Methodology, Writing - Review & Editing, Supervision, Project administration, Funding acquisition.

## Competing interests

The authors declare no competing interests.

## Notes

https://doi.org/10.5281/zenodo.15966533

https://doi.org/10.5281/zenodo.15926601

### Competing Interest Statement

The authors have declared no competing interest.

