## Supplementary Information for "Kinetic parameter prediction using neural networks identifies limitations to C_4_ photosynthesis"

### Supplementary Note 1

Identification of parameters that determine the feasibility of model simulations

To train a classifier to discriminate between feasible and infeasible parameter vectors, we considered two different definitions of feasibility: (1) the parameter set results in model simulations satisfying all criteria for physiologically relevant parameterizations defined above, and (2) the simulation results parameter set satisfy at least the technical criteria (1-3) of the definition. For each definition, ten subsets of the parameter samples were sampled, stratified according to the number of feasible and infeasible instances, each including n=472,392 and n=243,864 for the two definitions, respectively. First, the data was z-transformed and split into training (80%) and a test set (20%). The discrimination between feasible and infeasible parameter sets using decision trees has been successful for a kinetic model of *Escherichia coli* ^1^. Therefore, multiple ensemble classification models from scikit-learn (AdaBoostClassifier, BaggingClassifier, ExtraTreesClassifier, GradientBoostingClassifier, HistGradientBoostingClassifier, IsolationForest, RandomForestClassifier, Pedregosa *et al.*, 2011), tested with default settings on one of the sampled subsets, the histogram-based gradient boosting classification tree (HistGradientBoostingClassifier, HGBT) performed best considering accuracy, F1-score, and Matthews correlation coefficient for both definitions (Supplementary Fig. 15). The model hyperparameters were tuned by testing 100 parameter sets selected randomly from parameter distributions with five-fold cross-validation. As a result, the parameters yielding the best accuracy score (0.73 and 0.85 for definitions (1) and (2)) were the same for both definitions: l2_regularization=0.91, learning_rate=0.12, max_bins=183, max_depth=14, max_features=0.89, max_iter=177, max_leaf_nodes=74, and min_samples_leaf=71. After training the HGBT classifier, it showed an average accuracy score of 0.73 and 0.85 when applied on the unseen test set (precision=0.77 and 0.87, F1-score=0.72 and 0.84, recall=0.67 and 0.82, Matthews correlation coefficient=0.47 and 0.70). The importance of features in the model were determined by permuting each feature vector and comparing the resulting accuracy score to the performance with the unpermuted features (scikit-learn function permutation_importance). The reduction in accuracy score is then determined by the difference in accuracy between the original and the respective permuted feature sets. The calculation of feature importances for each feasibility definition was performed for each stratified subset and the highest average importances are shown in Supplementary Fig. 16. Partial dependences of the parameters were determined using the scikit-learn function PartialDependenceDisplay.

By inspecting the feature importance of the classifier that additionally considered biological feasibility, we found that the ten most important features for classification accuracy included parameters related to electron transport, RuBisCO (carboxylation), PEP carboxylase, sucrose synthesis, stomatal conductance, starch synthesis, and RuBisCO shunt (Supplementary Fig. 16A). The curvature parameter of the light response of the electron transport rate (Q32 or $\theta$ ^3,4^) showed a strong influence on the feasibility of a parameter vector (Supplementary Fig. 16A). In the training set of parameter samples, which contained equal numbers of feasible and infeasible parameter vectors, this parameter had an average value of 0.68 ± 0.19 (mean ± standard deviation), in line with its value of 0.7, commonly used in the FvCB model ^5^. From a theoretical point of view, this parameters should range between zero and one, to appropriately describe a non-rectangular hyperbola. Concordantly, we observed that in more than 93% of the parameter vectors with Q32 values greater than 1.0 were infeasible, demonstrating the importance of this parameter for the feasibility of the model. Further, we determined the partial dependences of features, given by their marginal effects on the classification outcome (Supplementary Fig. 17). The results show a clear threshold behavior for Q32 (Supplementary Fig. 17A), that could not be observed for features with lower importance.

When parameter feasibility was solely defined by the ability to simulate the model ODEs, only 12.2 % of the parameter vectors were infeasible. When the HGBT classification model was re-trained using this definition for model feasibility, the performance on the test set increased to an accuracy of 0.85, while the set of most important features remained largely the same (Supplementary Fig. 16B). However, here the most important parameters were all related to photosynthetic electron transport (Q32, X32, and Y32) and their impact on the accuracy of the model doubled compared to their effect in the classifier for the biologically-inspired definition of model feasibility.

### Supplementary Note 2

Identification of important parameters that determine the MSE of surrogate model predictions

To determine which parameter values lead to a reduced performance of the surrogate model, we trained a random forest regression model to determine parameters and potentially critical parameter values that determine the prediction error (MSE) of the surrogate model for A/CO_2_ and A/light curves. To address the inherent bias in the MSE values across the entire test set, we created a balanced subset, which contains 90 instances from each of five histogram bins. The random forest regression model was fitted to the entire subset, because the trained model was intended to describe the data and does not need to be generalized. The fitting process was repeated ten times, because we observed that the importance of features differed between different random number seeds. The trained regression model could replicate the training data with a coefficient of determination of R^2^=0.87 ± 0.01 (mean ± standard deviation) across the ten repetitions.

We found that the feature with the highest impact on the MSE of the surrogate model predictions was the V_max_ value of the chloroplastic Transketolase (Supplementary Fig. 18A). The partial dependence of the random forest regression model on this parameter shows that the average predicted MSE of the surrogate model predictions remains stable between 0.2 and 0.4 but increases for values outside this range (Supplementary Fig. 18B). An inspection of the CV of parameters associated with identical curve pairs revealed that the K_M_ values of the Transketolase for xylulose-5-phosphate and sedoheptulose-7-phosphate as well as its V_max_ value were among the parameters with the highest CV. In addition, the K_M_ value of Transketolase for glyceraldehyde-3-phosphate was among the most important features that determine the MSE of the surrogate model. These insights could guide future efforts aimed at improving surrogate model predictions which may include additional samples at the extreme ends of the distribution of the parameters associated with increased MSE values.

### Supplementary Note 3

Estimation of initial values for parameter sampling to generate the artificial dataset

- This section was adapted from Xu et al. 2025 ^6^ -

#### Gas exchange measurements

The experimental data for this study were obtained from the field trials of the maize MAGIC population ^7^, conducted at the National Institute of Agricultural Botany (NIAB, Cambridge, UK) over three consecutive seasons (2021, 2022, and 2023). The full details of the experimental design have previously been described in full detail ^8^. For this study, we made use of gas exchange measurements under varying ambient CO_2_ partial pressure and light intensities, which can be represented as A/CO_2_ and A/light curves. A/CO_2_ curves were available for 78, 88 and 91 recombinant inbred lines with A/CO_2_ curves in the three corresponding years, respectively. A/light curves were only available for 314 genotypes from 2022 and 2023, while photosynthetic rate at saturating light were recorded for 151 lines in 2021. The estimation of kinetic parameters was conducted for 68 genotypes with both A/CO_2_ and A/light curves from 2022 and 2023.

The net CO_2_ assimilation rate ($A_{net}$) and stomatal conductance ($g_{s}$) were measured at twelve ambient CO_2_ concentrations, starting at 400 µbar. Once $A_{net}$ stabilized, measurements were recorded every 120 s as the CO_2_ level increased to 600, 800, 1000, and finally up to 1250 µbar. Afterward, the CO_2_ partial pressure was restored to 400 µbar, followed by 300, 250, 200, 100, 75, and finally 25 µbar. Throughout A/CO_2_ measurements, the light intenstiy was kept constant at 1800 μmol m^-2^ s^-1^ and the exchanger temperature was set to 25 °C. The A/light curves were measured under a constant ambient CO_2_ (400 μbar) and temperature (25 °C). The initial light intensity level was set at 1800 μmol m^-2^ s^-1^ and then sequentially reduced to 1100, 500, 300, 150, and 50 μmol m^-2^ s^-1^.

#### Kinetic model of C_4_-photosynthesis

The C_4_-photosynthesis model used in this study included 84 metabolites whose concentration changes were determined by the 109 mass balances in the system, involving 123 reactions ^9^. The reaction rates involved 236 parameters, which we classified into four groups: (i) maximum velocities (V_max_), (ii) Michaelis-Menten constants (K_M_), (iii) activation rate constants of light-regulated enzymes, and (iv) the membrane permeability of specific metabolites. Since V_max_ values are the product of enzyme turnover number (k_cat_) and total enzyme concentration, this type of parameters can vary across different seasons. Thus, V_max_ parameters for 2022 and 2023 were treated as separate variables. All other parameters were required to be the same between the two years, in line with biophysical constraints. In addition, we observed discrepancies between the equilibrium constants used in the original mode ^9^ and those provided by eQuilibrator (Beber et al., 2022). Therefore, we refined 29 out of the 35 equilibrium constants accordingly, while ensuring the stability of the dynamic system.

#### Parameterization of genotype-specific kinetic models

The objective of model parameterization was to identify the set of kinetic parameters that results in simulated profiles ($A_{net}$ and $g_{s}$) as close as possible to the measured data. The standard distance metric used for fitting was the chi-square error ($\chi^{2}$), calculated using the formula:

$$\chi^{2}=\sum_{t} \frac{\left( O_{t}-S_{t} \right)^{2}}{\sigma_{t}^{2}},$$

(14)

where $O_{t}$ and $\sigma_{t}$ denote the measured data and their standard deviation at a given time point, $t$, while $S_{t}$ is the simulation result at the same time point. The advantage of using this metric is that it allows us to determine if the resulting fit is statistically significant at a given significance level, defined by the degrees of freedom, which corresponds to the number of data points minus the number of fit parameters.

Optimization of the non-linear objective ($\chi^{2}$), which embeds ordinary differential equation (ODE) simulations and involves a large number of parameters, does not guarantee to reach the global optimum. This is a common challenge in optimization problems and various algorithms have been developed to address it. In this study, we used PESTO’s parallel tempering method for parameterization ^10^. PESTO is a Bayesian approach that provides a probability distribution for the fitted parameters allowing the most probable value to be selected ^11^. Like other Bayesian methods, PESTO also provides confidence intervals for sampled parameters, enhancing the reliability of the parameter estimates. The classical Markov-chain Monte Carlo (MCMC) method evaluates the system’s energy using a single stochastic process and accepts or rejects updates based on the temperature, which is an auxiliary variable of the sampling approach. At high temperature, the system explores larger space, while at low temperature, the system may become trapped in local energy minima. Parallel tempering ^12^ was developed to address this issue by simulating replicas of the original system at different temperature and allowing exchange of complete configurations between systems, enabling systems at low temperature to escape local minima.

The required data for the optimization included:

1. Measured photosynthetic rates in response to environmental perturbations that reflect how a genotype responds to environmental changes:

- mean and standard deviation of measured photosynthetic rates (A/CO_2_) and stomatal conductances ($g_{s}$) across replicates at different ambient CO_2_ partial pressures.
- mean and standard deviation of photosynthetic rates (A/light) under varying light intensities

2. Initial guess of kinetic parameters ($\mathbf{k}_{\mathbf{0}}$), based on maize-specific parameters from ^9^, which were obtained from literature references, adapted from other kinetic models, or estimated from model simulation.
3. Initial metabolite concentrations ($\mathbf{x}_{\mathbf{t}_{\mathbf{0}}}$), required as an initial state of the ODE simulations.

We used the initial metabolite concentrations that used for simulation ^9^. Simulation of $A_{net}$ and $g_{s}$ was achieved by integrating the ODEs in the C_4_-photosynthesis model, given the initial metabolite concentrations, $\mathbf{x}_{\mathbf{t}_{\mathbf{0}}}$, sampled kinetic parameters ($\mathbf{k}_{\mathbf{sampled}}$) and environmental factors (CO_2_ and light) as inputs. The simulation starts with $\mathbf{x}_{\mathbf{t}_{\mathbf{0}}}$ and the first ambient CO2 level $CO_{2}(1)$, allowing the system to reach steady state, at which $A_{net}$ and $g_{s}$ were recorded as $\mathbf{ACasim}$(1) and $\mathbf{gssim}$(1), respectively. Subsequently, the ambient CO_2_ partial pressure was changed to $CO_{2}$(2) and the simulation was carried out for 120 s, to mimic observations, using the final metabolite concentrations from the first step. This process was repeated for the remaining measured CO_2_ levels, at constant mean chamber air temperature ($T_{air}$) for the given genotype. For simulation under changing light intensity, a similar procedure was followed, with a constant air temperature of 25 °C. The simulated A/CO_2_, $g_{s}$, and A/light curves were used to calculate the $\chi^{2}$ statistic, based on measured profiles (Eq. (14)). The curve simulation and calculation of the $\chi^{2}$ value was integrated as the objective for MCMC sampling.

The sampling algorithm started with the initial guess of kinetic parameters ($\mathbf{k}_{\mathbf{0}}$) and calculated the $\chi^{2}$ between measured and simulated profiles. If a new set of sampled parameters resulted in a smaller $\chi^{2}$, they were included in the ensemble; otherwise, they could be accepted if a random probability is below acceptance probability prescribed by the system temperature and posterior value. The sampling and updating of parameters were performed by parallel tempering algorithm. This process continued until a full ensemble of parameter sets was generated for each genotype.

### Supplementary Figures

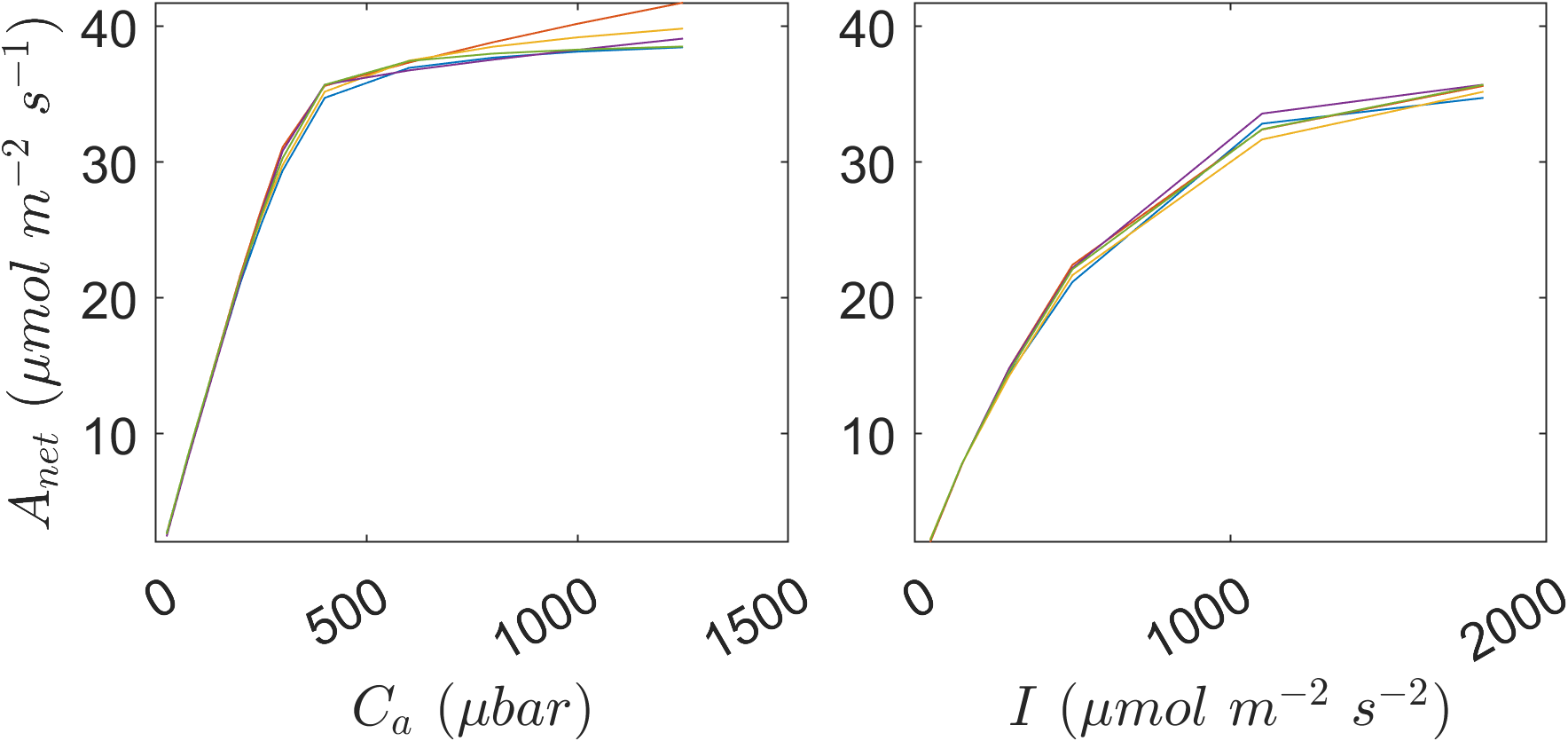

**Supplementary Figure 1. Example of parameter sets resulting in identical** $\boldsymbol{A}_{\boldsymbol{net}}$ **response curves.** These plots show A/CO_2_ and A/light curves simulated using five different parameter sets, which were considered identical for the assessment of parameter identifiability.

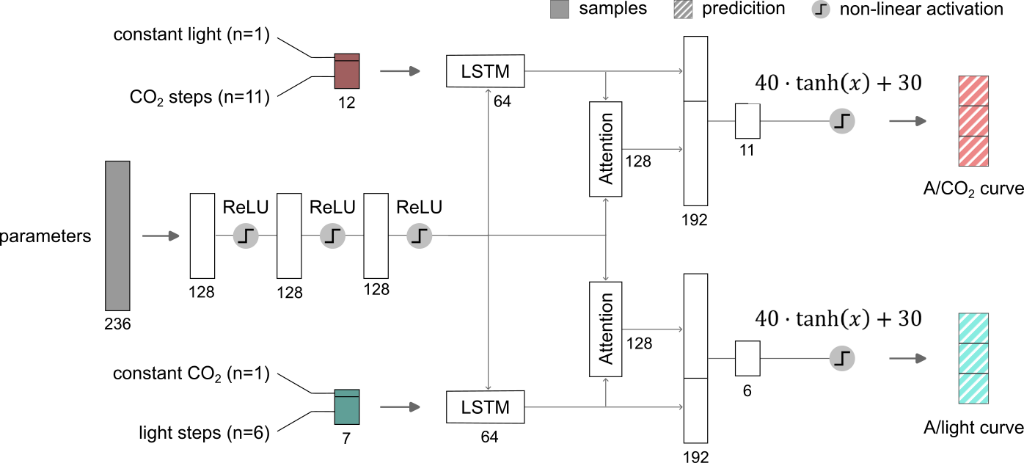

**Supplementary Figure 2. Detailed architecture of the surrogate neural network model.** A neural network was trained to predict A/CO_2_ and A/light curves from environmental inputs and a parameter vector, to serve as a surrogate model for the ODE model. The empty rectangles denote linear layers, which are connected by non-linear activation functions. The different layers are connected by arrows indicating the forward direction of the network. The numbers below the layers indicate their output dimension.

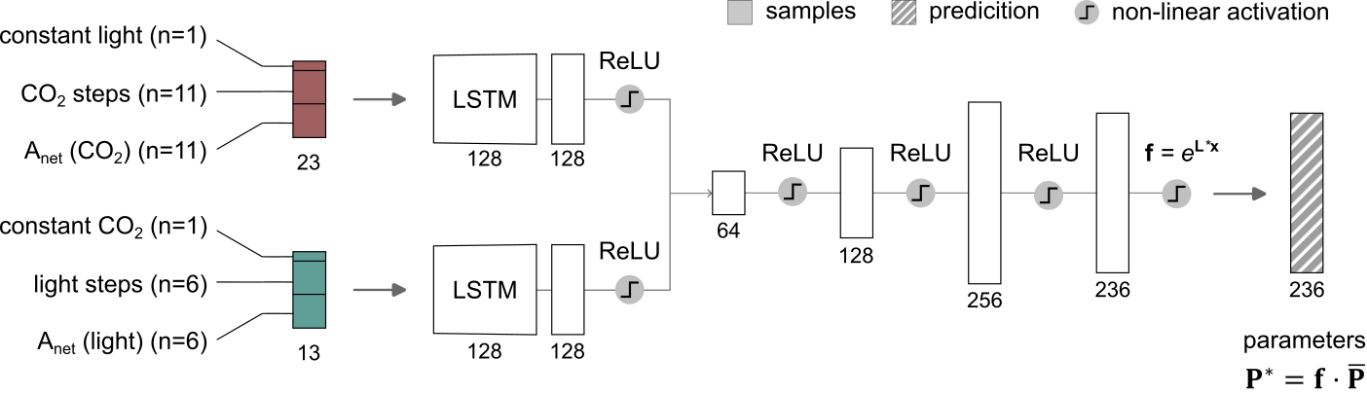

**Supplementary Figure 3. Detailed architecture of the C4TUNE model.** A neural network was trained to predict parameter vector from A/CO_2_ and A/light curves and associated environmental inputs. The empty rectangles denote linear layers, which are connected by non-linear activation functions. The different layers are connected by arrows indicating the forward direction of the network. The numbers below the layers indicate their output dimension. $\mathbf{f}$: predicted parameter deviations, $\mathbf{x}$ output values from the last linear layer of the network, $\mathbf{L}$: Cholesky decomposition of the covariance matrix of log-transformed parameters from the training or test set, $\mathbf{P}^{\mathbf{*}}$: parameter predictions, $\bar{\mathbf{P}}$ average parameters of the training or test set.

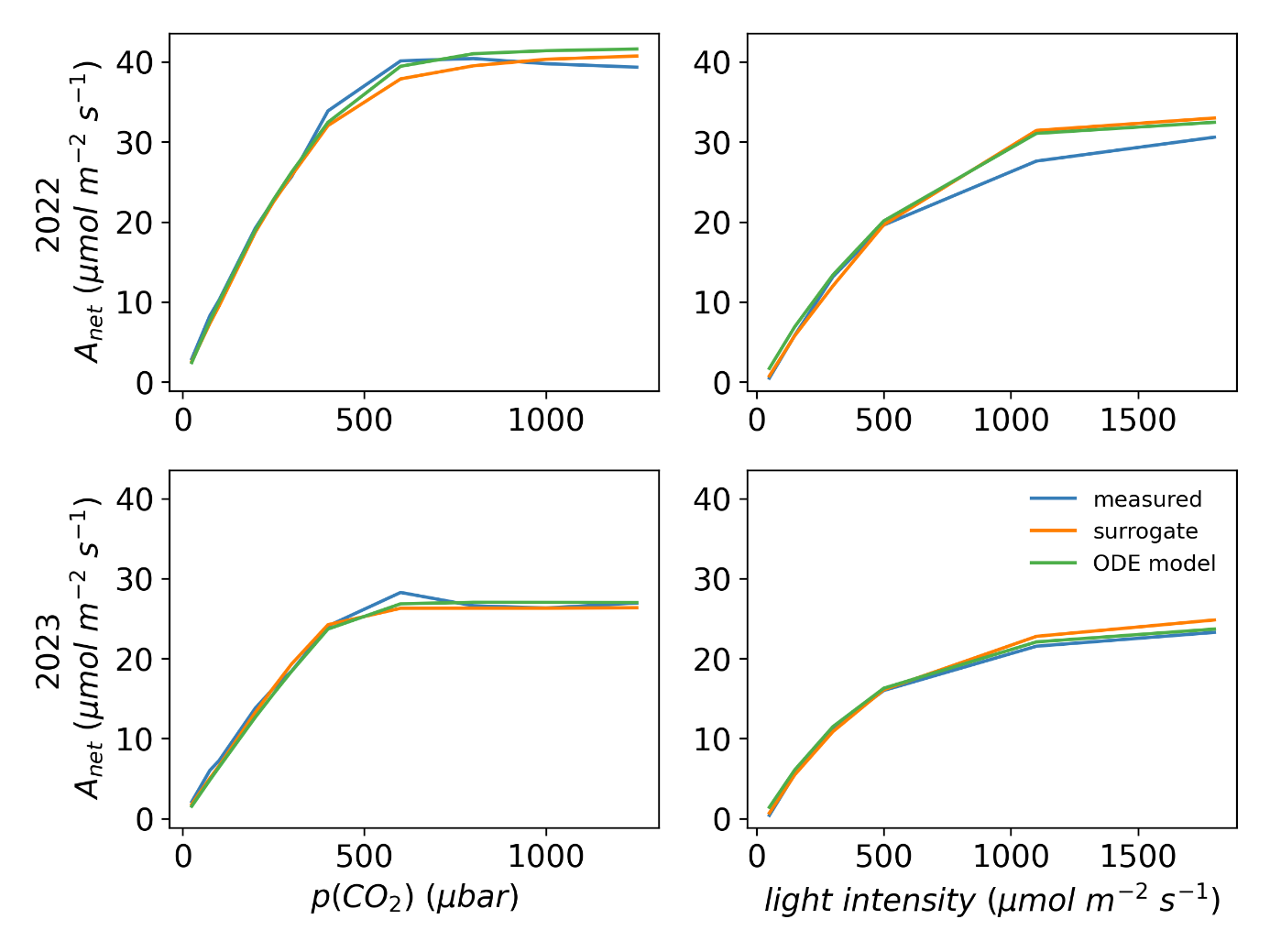

**Supplementary Figure 4. Example curve simulations based on predicted parameters using C4TUNE.** Gas exchange measurements (A/CO_2_ and A/light curves) for one of 68 maize accessions in two growing seasons (2022 and 2023) were randomly selected and used as inputs for C4TUNE to predict the associated ODE model parameters. The line graphs show the experimentally measured response in net assimilation rate ($A_{net}$) to ambient CO_2_ partial pressure, $p\left( CO_{2} \right)$, and light intensity as well as corresponding model simulations using the surrogate model and the ODE model ^9^.

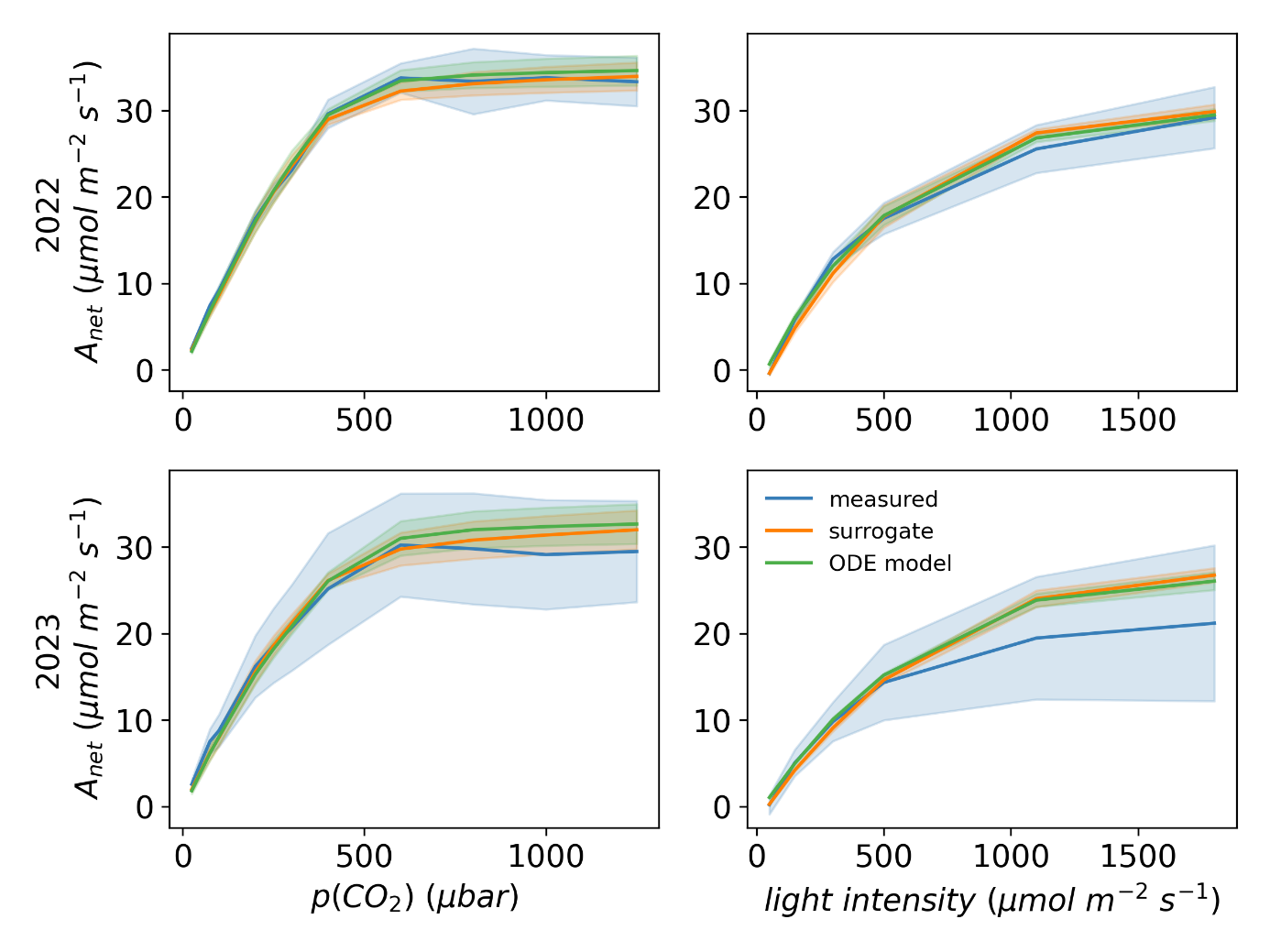

**Supplementary Figure 5. Example curve simulations based on predicted parameters for sampled** $\boldsymbol{A}_{\boldsymbol{net}}$ **response curves within experimental error using C4TUNE.** Gas exchange measurements (A/CO_2_ and A/light curves) for one of 68 maize accessions in two growing seasons (2022 and 2023) were randomly selected, and random curves were sampled within the experimental error at each step of the respective curves for the respective genotype and year. The sampled curves were used as inputs for C4TUNE to predict the associated ODE model parameters. The lines show the experimentally measured responses in net assimilation rate ($A_{net}$) to ambient CO_2_ partial pressure, $p\left( CO_{2} \right)$, and light intensity as well as corresponding model simulations using the surrogate model and the ODE model ^9^. The shaded areas show the standard deviation.

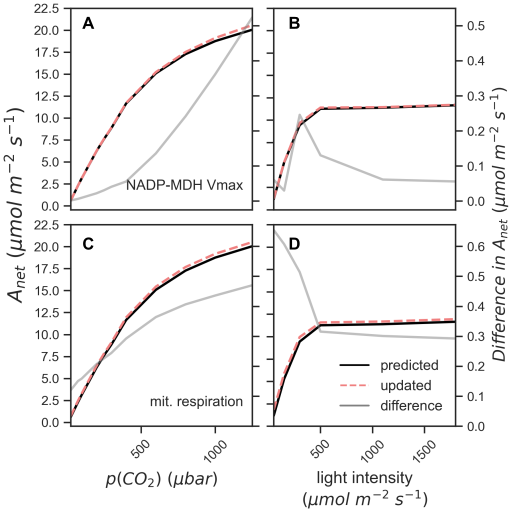

**Supplementary Figure 6. Simulation results after optimization of selected model parameters.** The simulations were carried out for accession SSA_00006, which showed the lowest median $A_{net}$ values across all CO_2_ and light steps in the 2022 growth period. The selected parameters were optimized by replacing them with the extreme values of the predicted parameters across all 68 genotypes, depending on the sign of their correlation with $A_{net}$. The plots show the comparison the simulated $A_{net}$ response curves with the predicted (“predicted”) and the updated parameters (“updated”). **(A, B)** selected target from correlations between parameter values and $A_{net}$ values in A/CO_2_ curves, **(C, D)** selected target from correlations between parameter values and $A_{net}$ values in A/light curves. Out of all identified targets, these two were selected because they showed the highest increase in $A_{net}$, respectively.

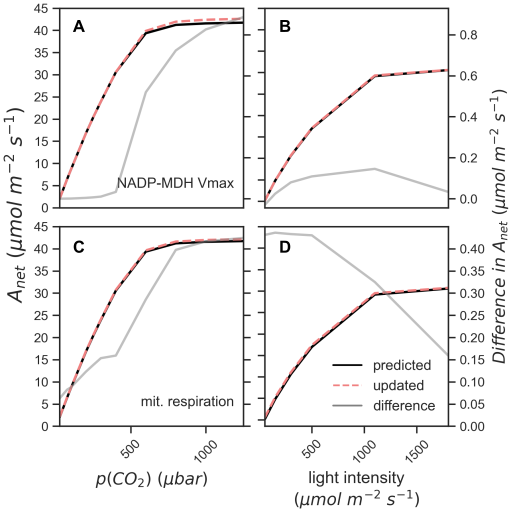

**Supplementary Figure 7. Simulation results after optimization of selected model parameters.** The simulations were carried out for accession SSA_00001, which showed an average median $A_{net}$ values across all CO_2_ and light steps in the 2022 growth period. The selected parameters were optimized by replacing them with the extreme values of the predicted parameters across all 68 genotypes, depending on the sign of their correlation with $A_{net}$. The plots show the comparison the simulated $A_{net}$ response curves with the predicted (“predicted”) and the updated parameters (“updated”). **(A, B)** selected target from correlations between parameter values and $A_{net}$ values in A/CO_2_ curves, **(C, D)** selected target from correlations between parameter values and $A_{net}$ values in A/light curves. Out of all identified targets, these two were selected because they showed the highest increase in $A_{net}$, respectively.

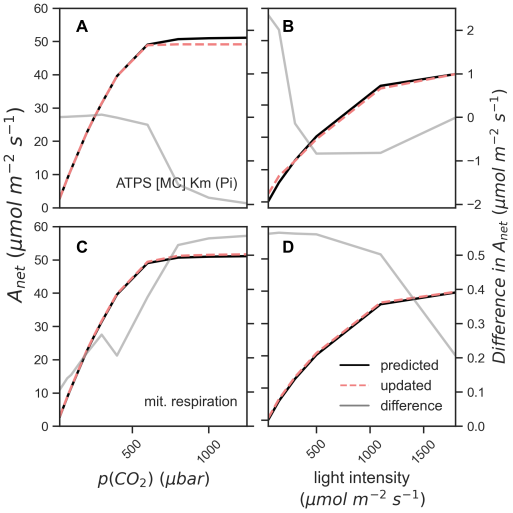

**Supplementary Figure 8. Simulation results after optimization of selected model parameters.** The simulations were carried out for accession SSA_00367, which showed the highest median $A_{net}$ values across all CO_2_ and light steps in the 2022 growth period. The selected parameters were optimized by replacing them with the extreme values of the predicted parameters across all 68 genotypes, depending on the sign of their correlation with $A_{net}$. The plots show the comparison the simulated $A_{net}$ response curves with the predicted (“predicted”) and the updated parameters (“updated”). **(A, B)** selected target from correlations between parameter values and $A_{net}$ values in A/CO_2_ curves, **(C, D)** selected target from correlations between parameter values and $A_{net}$ values in A/light curves. Out of all identified targets, these two were selected because they showed the highest increase in $A_{net}$, respectively.

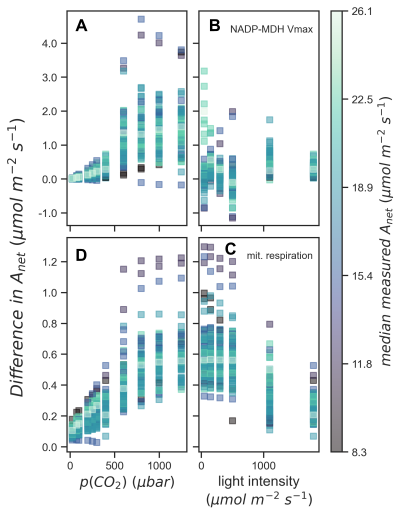

**Supplementary Figure 9. Simulation results after optimization of selected model parameters across all genotypes.** The simulations were carried out for all 68 maize genotypes, based on the parameters predicted for the 2022 growth period. The selected parameters were optimized by replacing them with the extreme values of the predicted parameters across artificial dataset, depending on the sign of their correlation with $A_{net}$. The plots show the difference between the simulated $A_{net}$ values with predicted and optimized parameter values. The color code indicates the median of the measured $A_{net}$ values per accession. **(A, B)** selected target from correlations between parameter values and $A_{net}$ values in A/CO_2_ curves, **(C, D)** selected target from correlations between parameter values and $A_{net}$ values in A/light curves. Out of all identified targets, these two were selected because they showed the highest median increase in $A_{net}$ across all accessions, respectively.

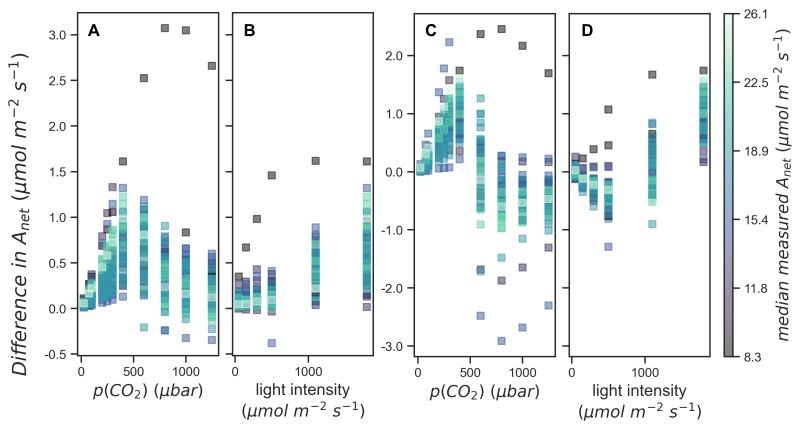

**Supplementary Figure 10. Simulation results with increased V_max_ values for the RuBisCO carboxylation and oxygenation reactions.** The simulations were carried out for all 68 maize genotypes, based on the parameters predicted for the 2022 growth period. To simulate an increased RuBisCO content, the V_max_ values for the RuBisCO carboxylation and oxygenation reactions were increased by replacing it with the maximum value of the predicted parameters across the predicted parameters for all 68 accessions (**A, B**) or across the artificial dataset (**C, D**). The plots show the difference between the simulated $A_{net}$ values with predicted and optimized parameter values. The color code indicates the median of the measured $A_{net}$ values per accession.

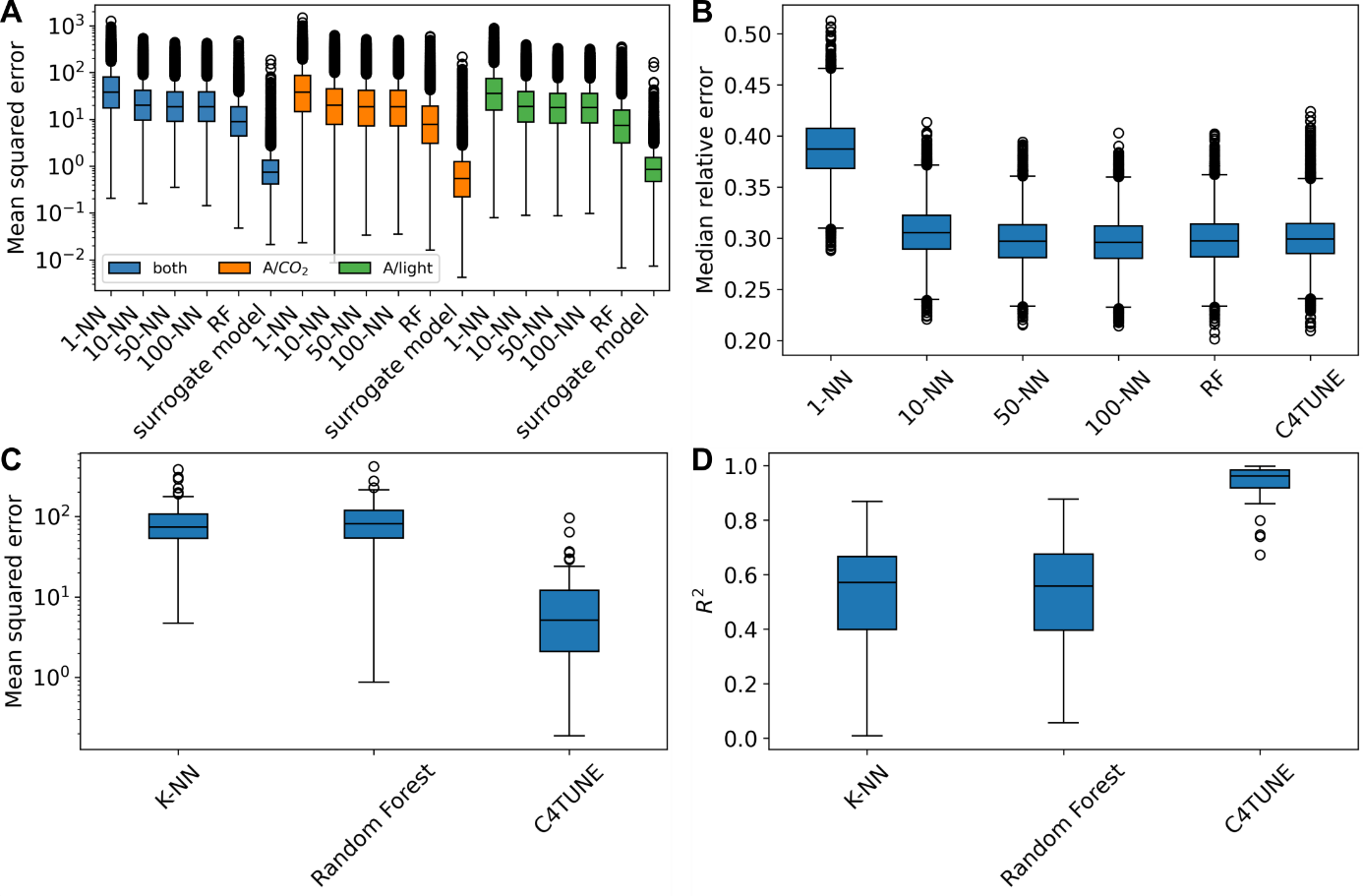

**Supplementary Figure 11. Comparison of prediction performances between C4TUNE and classical machine learning approaches.** A subset of 10^5^ parameters and associated ODE model simulations of A/CO_2_ and A/light curves was divided into a training (80%) and a test (20%) set to train K-nearest neighbors (K-NN) and random forest regression models. The models were trained to predict both curve types individually and jointly based on z-transformed parameter values. Further, K-NN ($K\in\left\{ 1, 10, 50, 100 \right\}$) and random forest regression models were trained to predict deviations from average parameter values, which is the same endpoint as in C4TUNE, from z-transformed $A_{net}$ response curves. The predicted parameters were used to simulate A/CO_2_ and A/light curves using the ODE model. **A.** Mean squared error between sampled and predicted curves for the different regression models for either joint (“both”) or individual curve type prediction. The results for the surrogate model trained using the full dataset is shown as a comparison. The performance of all approaches was assessed using the test set. **B.** Median relative errors between predicted and sampled parameters. The results of C4TUNE are shown as a comparison, which was trained on the full dataset. However, the Cholesky decomposition was determined based on the test set. The performance of all approaches was assessed using the test set. **C., D.** Mean squared errors and coefficients of determination (R^2^) between curves in the test set and simulated using the ODE based on the predicted parameters (n=100). For the R^2^ values, only values between 0 and 1 are shown. The numbers of negative R^2^ values were 39 (K-NN, K=100), 42 (random forest), and 5 (C4TUNE).

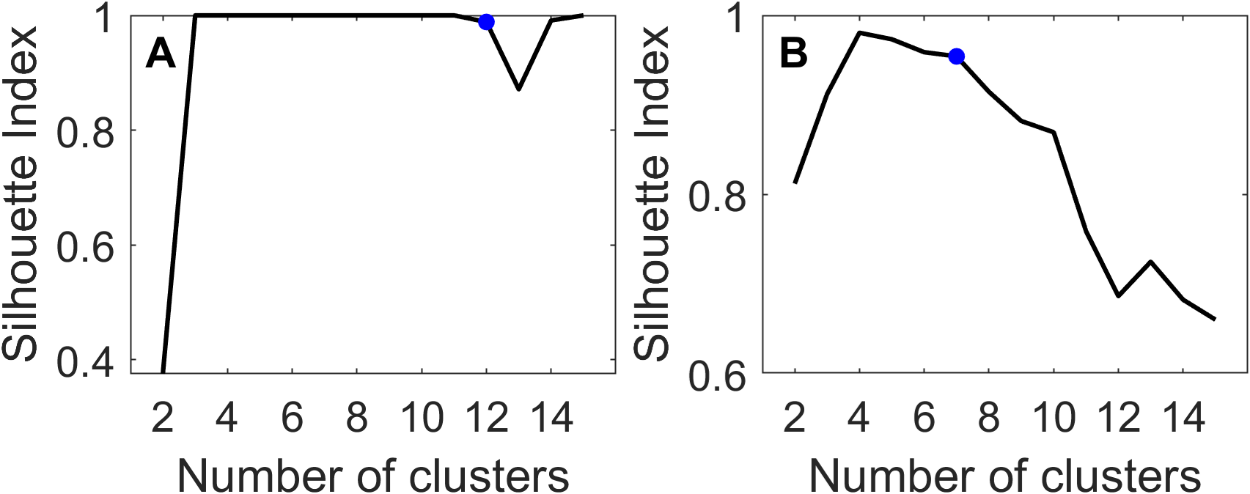

**Supplementary Figure 12. Clustering performance of A/CO_2_ and A/light curves.** A subsample of the generated training dataset (n=10,000) was clustered based on the responses of $A_{net}$ to different levels of CO_2_ **(A)** and light **(B)**. The clustering was performed by transforming the data by non-linear embedding and applying spectral clustering with different numbers of clusters (K). The median Silhouette Index was calculated for each K. The blue dot indicates the K that was chosen for the final clustering.

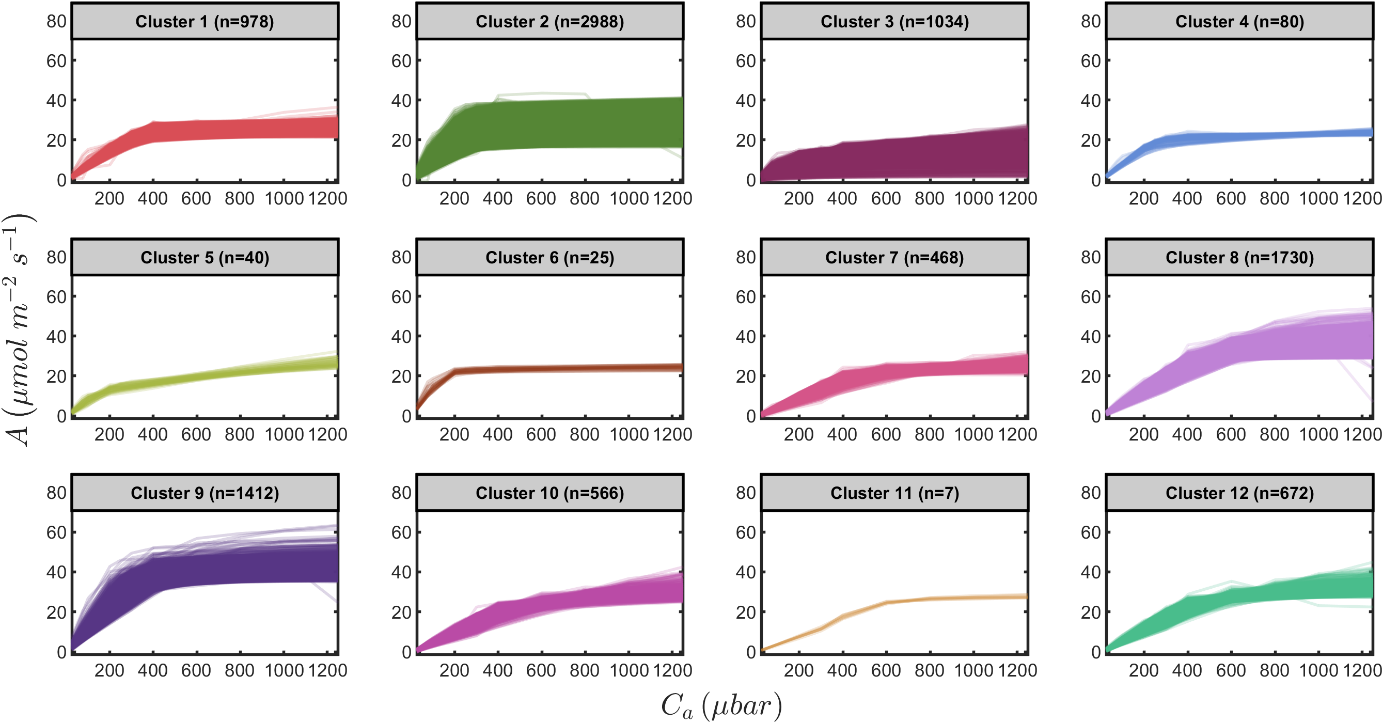

**Supplementary Figure 13. Clustering of A/CO_2_ curves.** A subsample of the generated training dataset (n=10,000) was clustered based on the responses of $A_{net}$ to different levels of CO_2_. The clustering was performed by transforming the data by non-linear embedding and applying spectral clustering (Methods).

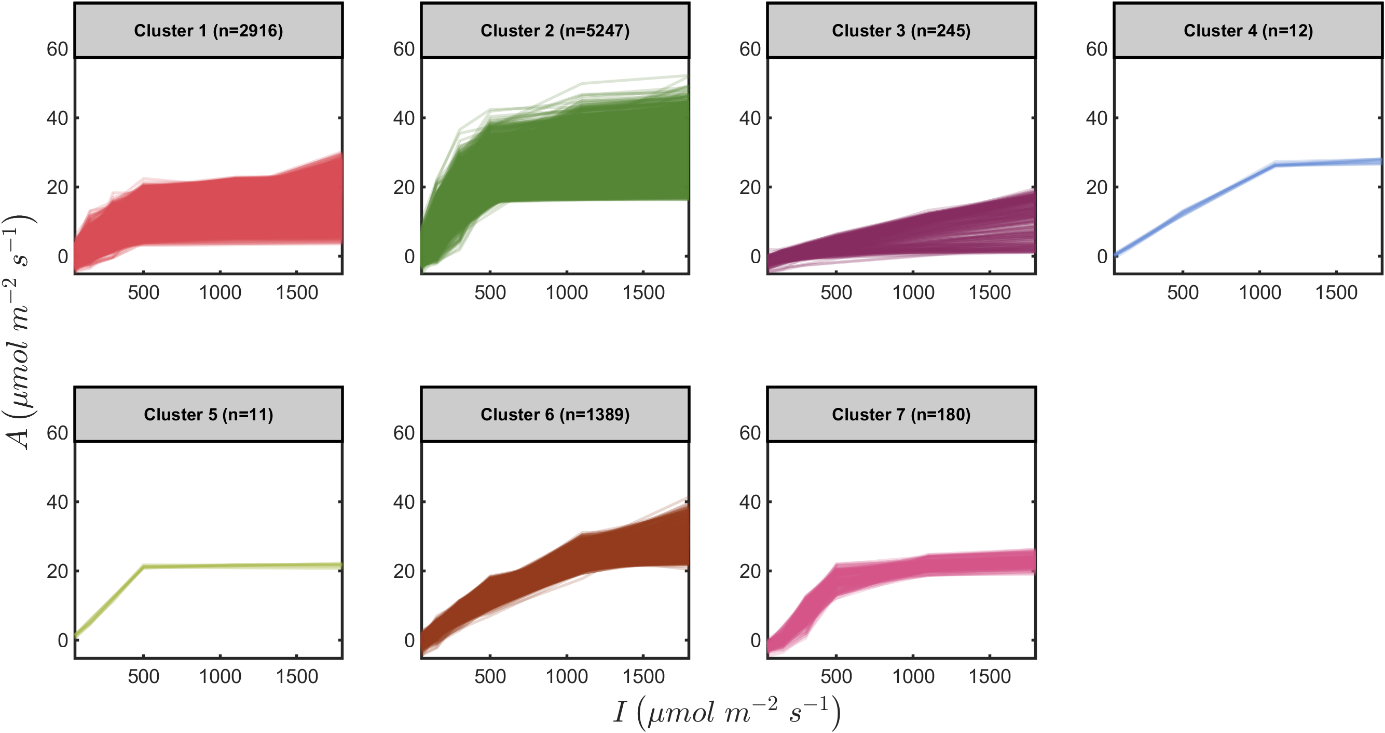

**Supplementary Figure 14. Clustering of A/light curves.** A subsample of the generated training dataset (n=10,000) was clustered based on the responses of $A_{net}$ to different levels of light intensity. The clustering was performed by transforming the data by non-linear embedding and applying spectral clustering (Methods).

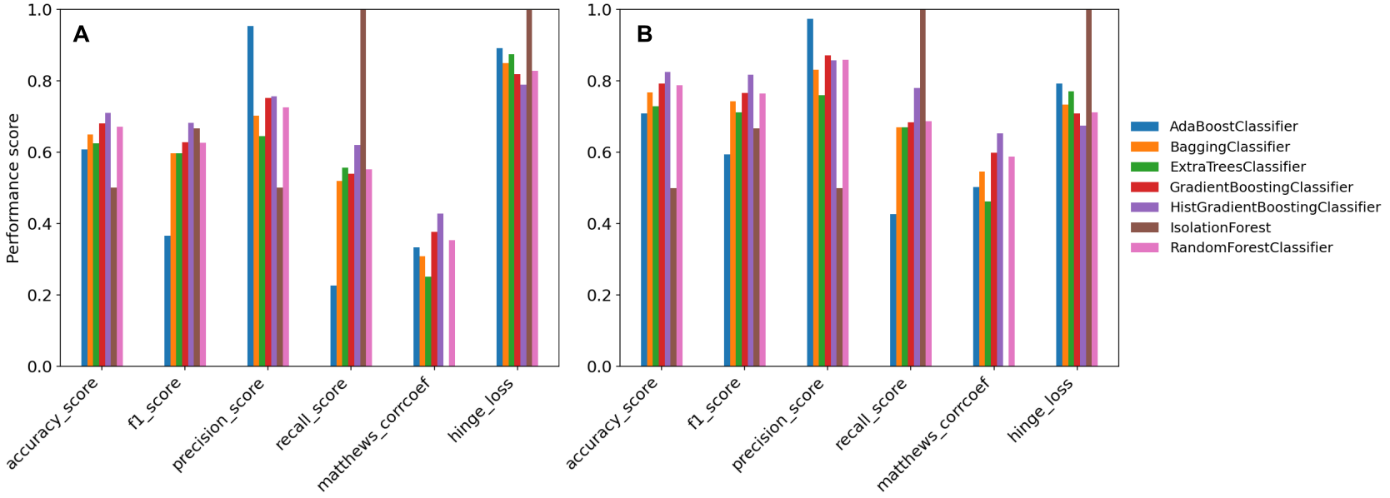

**Supplementary Figure 15. Default performance scores of different classifier models for the prediction of infeasible parameter sets.** Different classification models in sklearn ^2^ were trained with default settings classify parameter vectors as feasible if the resulting model can be simulated (considered in **A** and **B**) and the resulting $A_{net}$ response curves are biologically relevant (only considered in A); otherwise, a parameter vector is considered infeasible. The models were trained on balanced datasets with respect to the numbers of feasible and infeasible parameter vectors, depending on the definition of feasibility. The data z-transformed and divided into a training (80%) and a test set (20%), respectively. The performance scores shown were obtained by applying the trained models to the test set.

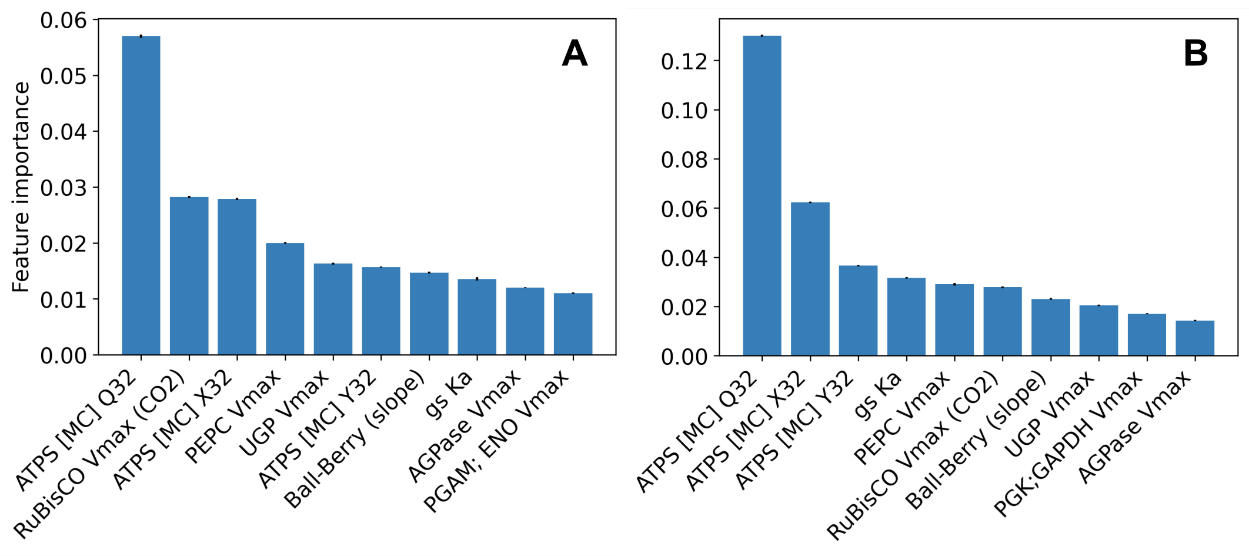

**Supplementary Figure 16. Most important features for the classification of parameter vectors based on feasibility.** Histogram-based gradient boosting classification trees (HGBT) were trained to classify parameter vectors as feasible if the resulting model can be simulated (considered in **A** and **B**) and the resulting $A_{net}$ response curves are biologically relevant (only considered in A); otherwise, a parameter vector is considered infeasible. The model training was repeated over ten subsets of the parameter samples, stratified according to the number of feasible and infeasible instances. The feature importances were determined by random permutation of the feature vectors and scoring the change in accuracy to the original classifier in a five-fold cross validation per subset. The bars show the average feature importances and the error bars show the standard error of the mean across the ten subsets. ATPS: ATP-synthase, PEPC: phosphoenolpyruvate carboxylase, UGP: UTP---glucose-1-phosphate uridylyltransferase, AGPase: ADP-glucose pyrophosphorylase, PGAM: phosphoglycerate mutase, ENO: 2-phosphoglycerate enolase, Q32: curvature parameter ($\theta$) for the response of electron transport to light, X32: light partition coefficient, Y32: J_max_ partition coefficient, gs: stomatal conductance, MC: mesophyll cell, Vmax: maximum reaction velocity, Ka: activation constant.

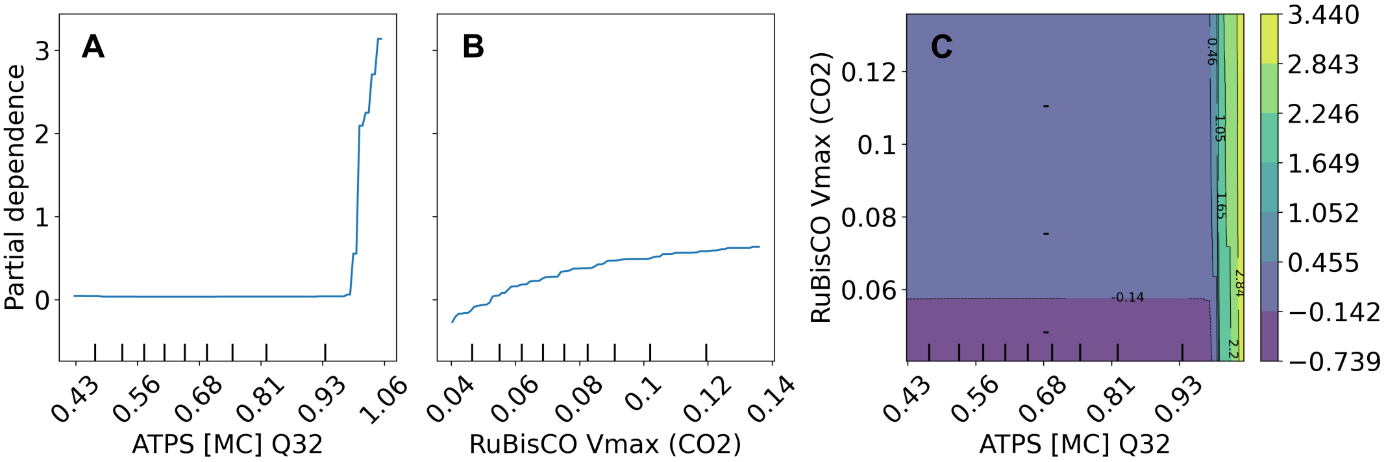

**Supplementary Figure 17. Partial dependences of the most important features for the classification of parameter vectors based on feasibility.** A histogram-based gradient boosting classification tree (HGBT) was trained to classify a parameter vector as feasible if the resulting model can be simulated and the resulting $A_{net}$ response curves are biologically relevant; otherwise, a parameter vector is considered infeasible. The individual partial dependences of the two most important features are shown, which show their marginal effects on the classification response **(A, B)** along the joint dependence of the classification result in the parameter values **(C)**. The colors in (C) depict the raw score of the HGBT classification model, where a score > 0.5 corresponds to the infeasible class. ATPS: ATP-synthase, Q32: curvature parameter ($\theta$) for the response of electron transport to light.

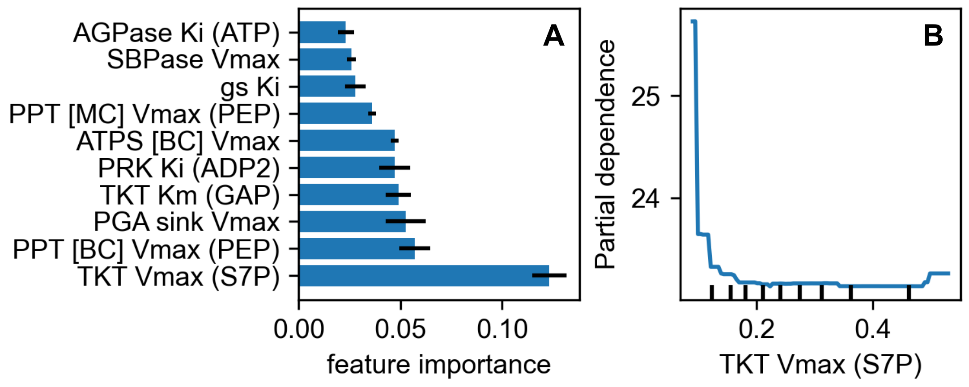

**Supplementary Figure 18. Important features for the surrogate model predictions.** A random forest regression model was trained to predict MSE values of the surrogate model predictions using associate	d parameter vectors was input. **A.** most important features for the R^2^ value of the model based on Gini importance. The bars show average values from ten repetitions, and the error bars represent the standard deviation. **B.** partial dependence of the regression output on the most important feature, i.e., its marginal effect on the predicted MSE values. AGPase: ADP-glucose pyrophosphorylase, SBPase: sedoheptulose-bisphosphatase, gs: stomatal conductance, PPT: PEP/phosphate transporter, PRK: phosphoribulokinase, TKT: transketolase; PEP: phosphoenolpyruvate, GAP: glyceraldehyde-3-phosphate, S7P: sedoheptulose-7-phosphate, V_max_: maximum reaction velocity, Km: Michaelis-Menten constant, Ki: inhibitory constant, the specificity of the constants is indicated in parentheses, MC: mesophyll cell, BC: bundle sheath cell.
